# NKCC1 modulates microglial phenotype, cerebral inflammatory responses and brain injury in a cell-autonomous manner

**DOI:** 10.1101/2021.01.21.427597

**Authors:** Krisztina Tóth, Nikolett Lénárt, Péter Berki, Rebeka Fekete, Eszter Szabadits, Balázs Pósfai, Csaba Cserép, Ahmad Alatshan, Szilvia Benkő, Dániel Kiss, Christian A. Hübner, Attila Gulyás, Kai Kaila, Zsuzsanna Környei, Ádám Dénes

## Abstract

The NKCC1 ion transporter contributes to the pathophysiology of common neurological disorders, but its function in microglia, the main inflammatory cells of the brain, has remained unclear to date. Therefore, we generated a novel transgenic mouse line in which microglial NKCC1 was deleted. We show that microglial NKCC1 shapes both baseline and reactive microglia morphology, process recruitment to the site of injury, and adaptation to osmotic stress in a cell-autonomous manner via regulating membrane potential and chloride fluxes. In addition, microglial NKCC1 deficiency results in increased expression of the D subunit of volume regulated anion channel (VRAC), NLRP3 inflammasome priming and production of interleukin-1β (IL-1β), rendering microglia prone to exaggerated inflammatory responses. In line with this, central (intracortical) administration of the NKCC1 blocker, bumetanide, potentiated intracortical lipopolysaccharide (LPS)-induced cytokine levels, whereas systemic bumetanide application decreased inflammation in the brain. Microglial NKCC1 KO animals exposed to experimental stroke showed significantly increased brain injury, inflammation, cerebral edema and worse neurological outcome. Thus, NKCC1 emerges as an important player in controlling microglial ion homeostasis and inflammatory responses through which microglia modulate brain injury. The contribution of microglia to central NKCC1 actions is likely to be relevant for common neurological disorders.

## Introduction

Members of the plasmalemmal cation-chloride cotransporter (CCC) family, such as the neuron-specific K^+^/Cl^-^ extruder, KCC2, and the ubiquitously expressed Na^+^-K^+^-2Cl^-^ cotransporter, NKCC1 (coded by the *Slc12a2* gene), have received a steeply increasing amount of attention in research on CNS diseases, ranging from neuropsychiatric diseases to epilepsy, stroke and dementia [1–6]. Notably, there’s an abundance of studies, which have shown that the mRNA and protein expression levels as well as the functionality of NKCC1 are enhanced in injured and post-traumatic neurons [7, 8]. This, in turn, has raised the obvious possibility that assuming a pathophysiological role for NKCC1, therapeutic actions might be achieved by pharmacological inhibition of this transporter. Indeed, data in a number of experimental studies based on systemic application of the selective NKCC1 blocker, the loop diuretic bumetanide, have suggested that this drug has therapeutic actions in diverse neuropathological conditions. These include neonatal seizures, temporal lobe epilepsy, autism spectrum disorders, schizophrenia, and brain edema after traumatic or hypoxic/ischaemic injury [1–4,7–12]. In most of these studies, bumetanide has been suggested to exert its therapeutic effects by acting directly on NKCC1 expressed in central neurons.

While bumetanide is routinely used to block neuronal NKCC1 in experiments performed *in vitro* [3,8,12], the view that the drug might have direct effects on central neurons *in vivo* has raised numerous questions. First, the poor pharmacokinetic properties of bumetanide, including a low permeability to active coupled extrusion across the blood-brain barrier (BBB) would imply that the drug does not reach pharmacologically relevant concentrations in the brain parenchyma, a prediction that has been experimentally verified [10,13,14]. Another important point in the present context is that NKCC1 is expressed within the immature and mature CNS mainly in non-neuronal cell types [15], such as oligodendrocytes and their precursors [16, 17], ependymal cells and astrocytes [18–21]. Thus, even if applied directly into brain tissue, bumetanide or any other NKCC1 blockers would be expected to inhibit this ion transporter in a wide variety of cell types.

Interestingly, common neurological disorders including those with altered NKCC1 activity [6,7,22] display broad neuroinflammatory changes. While the impact of NKCC1 function on inflammatory cytokine production has not been widely studied, emerging evidence indicates upregulated NKCC1 expression in response to inflammatory stimuli (LPS, IL-1β, TNF-α) [23–26]. Notably, while microglia are the main inflammatory cell type in the CNS, there is no information about the functional role of NKCC1 in microglia and whether microglial NKCC1 could provide a significant contribution to central NKCC1 actions under inflammatory conditions or after brain injury.

In the healthy brain, microglia regulate neuronal activity and responses, whereas altered microglial activity is linked with the pathophysiology of most common brain disorders, such as neurodegenerative and psychiatric diseases, stroke and epilepsy [27–33]. Extracellular accumulation of potassium is related to diverse neuropathological alterations [32, 34], and the contribution of microglial potassium channels and transporters to microglial activity is widely recognized. These include membrane-expressed potassium channels (Kv1.3, THIK-1, Kir2.1) that regulate microglial motility, immune surveillance and cytokine release, among others [35, 36]. Moreover, changes in intracellular potassium and chloride levels are associated with inflammasome activation contributing to the regulation of IL-1β release [37–39]. Interestingly, recent transcriptomic data verified high-level NKCC1 expression in microglia [21, 40]. However, currently no experimental data are available on the function of microglial NKCC1 under physiological and pathological conditions. To this end, we studied whether central and systemic NKCC1 blockade impact on central inflammatory changes differently and tested the hypothesis that NKCC1 is involved in the regulation of microglial inflammatory mediator production, neuroinflammation and brain injury.

## Results

### Systemic and central blockade of NKCC1 regulate LPS-induced inflammatory cytokine production in the brain in an opposite manner

To investigate whether *systemic blockade of NKCC1 actions* by bumetanide could alter inflammatory responses in the CNS, we injected mice either intraperitoneally or intracortically with bacterial lipopolysaccharide (LPS), while NKCC1 actions were blocked by systemic (intraperitoneal; ip.) bumetanide administration (Figure 1A). As expected, LPS administration by ip. injection triggered marked inflammation in the spleen and the liver (Supplementary Figure 1A), but caused low cytokine production in cortical brain tissues, which was not influenced by ip. bumetanide treatment (Figure 1A). However, bumetanide resulted in a small, but significant increase in LPS-induced IL-1β production in the spleen (Supplementary Figure 1A). In contrast, intracortical LPS administration triggered a robust inflammatory response in the brain as seen earlier [41], which was reduced by ip. bumetanide treatment. This was demonstrated by lower G-CSF, KC, IL-1β, and IL-1α levels in the brain (by 39.8%, 43%, 54.6% and 41%, respectively), while systemic cytokine levels were not altered (Supplementary Figure 1A).

**Figure 1.**
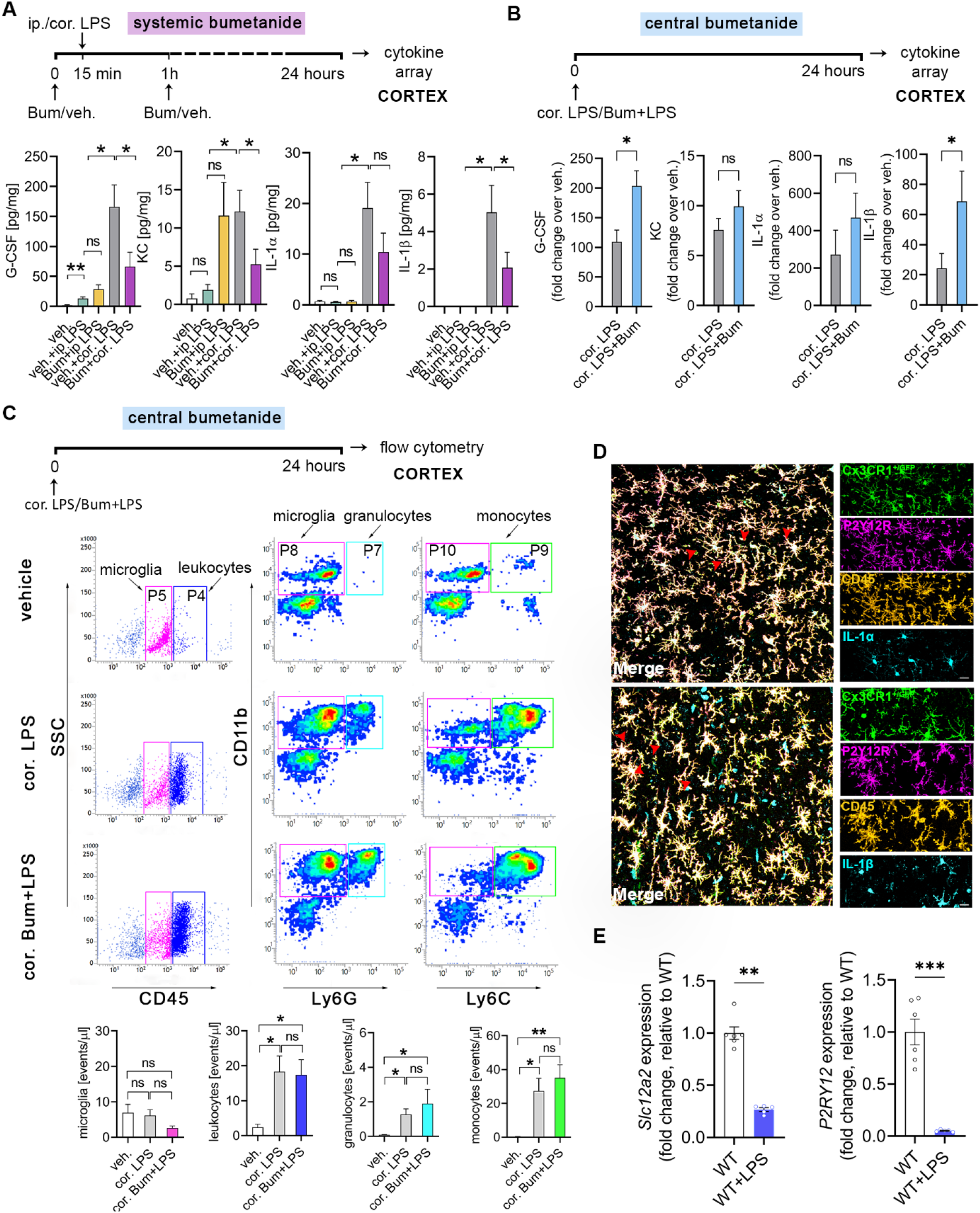
Systemic and central (intracortical) blockade of NKCC1 regulate LPS-induced inflammatory cytokine production in the brain in an opposite manner. **A**: Mice were subjected to either intraperitoneal (ip.) or intracortical (cor.) LPS injections, while NKCC1 was blocked by ip. bumetanide (Bum) administration. Central LPS injection triggers high cytokine (G-CSF, IL-1α, IL-1β) and chemokine (KC) responses in the brain compared to ip. LPS injection, which is blocked by ip. Bum administration. **B**: Central NKCC1 inhibition by intracortical Bum administration significantly increases GCSF and IL-1β levels. See also Supplementary Figure 1 for effects of systemic vs. central blockade of NKCC1 on LPS-induced cytokine responses in the periphery. **C**: Flow cytometric dot plots show that cortical administration of Bum does not affect the number of microglia (CD45^int^/P5 gate), and recruitment of leukocytes (CD45^high^/P4 gate), including monocytes (CD11b^+^, Ly6C^high^ /P9 gate), and granulocytes (CD11b^+^, Ly6G^high^/P7 gate) upon central LPS injection. **D**: The main source of IL-1α and IL-1β in the brain are microglia cells. Confocal images of Cx3CR1^+/GFP^ brain slices show IL-1α-CD45-P2Y12R (above, red arrowheads) and IL-1β-CD45-P2Y12R (below, red arrowheads) labelled cells after cortical LPS injection-induced inflammation. All data are expressed as mean±SEM. **E**: NKCC1 (encoded by *Slc12a2*) and P2Y12R gene expression is downregulated in microglia isolated from adult mice 24 hours after cisterna magna LPS application. **A**: One-way ANOVA followed by Tukey’s multiple comparison test; \**p*<0.05; N=6/group; **B**: Unpaired t-test; \**p*<0.05; N=9/group; Data were pooled from two independent studies. **C**: One-way ANOVA followed by Tukey’s multiple comparison test; \**p*<0.05; N=4/group. **D**: Scale: 25 μm; **E**: Unpaired t-test; \*\**p*<0.01, \*\*\**p*<0.001; N (WT)=6, N (KO)=5; Abbreviations: veh.: vehicle; ip: intraperitoneal; cor.: cortical; Bum: bumetanide; ns: not significant

To compare the effects of systemic versus central NKCC1 blockade by bumetanide, we next investigated the impact of *central (intracortical) bumetanide administration* on LPS-induced cytokine responses, by co-injecting bumetanide with LPS (Figure 1B) into the cerebral cortex. As expected, intracortical LPS without bumetanide resulted in a 20-50-fold increase in inflammatory cytokines/chemokines in the cerebral cortex compared to vehicle. Surprisingly, on top of this, bumetanide markedly potentiated LPS-induced G-CSF, KC, IL-1β and IL-1α levels in the brain (by 86.1%, 31.2%, 82.5% and 72.4%, respectively), in sharp contrast with the effects of ip. bumetanide treatment (Figure 1A). Intracortical bumetanide had no effect on systemic cytokine levels (Supplementary Figure 1B).

We next tested whether the effects of central bumetanide administration on increasing cytokine production might be explained by altered LPS-induced recruitment of leukocytes. However, while flow cytometry revealed increased number of infiltrating CD45^high^ leukocytes, in particular CD11b^+^ Ly6G^high^ granulocytes, and CD11b^+^ Ly6G^-^ Ly6C^high^ monocytes in response to central LPS injection compared to vehicle administration (Figure 1C), this was not altered by central bumetanide. The number of CD45^int^ CD11b^+^ microglia was not affected either (Figure 1C), indicating that the increased LPS-induced cytokine production observed upon central bumetanide treatment is not due to altered inflammatory cell numbers in the cerebral cortex. Unbiased densitometric analysis revealed that GFAP and AQP4 immunopositivity 24 hours after intracortical LPS injection were not altered by central bumetanide treatment (Supplementary Figure 2), suggesting that while bumetanide may also act on astroglial NKCC1 [18–20], marked changes in astrocyte phenotypes or perivascular astrocyte endfeet are unlikely to explain the effect of bumetanide on central inflammatory responses in the present study.

Microglia are the primary source of inflammatory cytokines in a number of neuropathologies, and IL-1β, also produced by microglia, is a key proinflammatory cytokine regulating the brain’s cytokine network [42, 43]. To investigate the cellular source of IL-1β in this experimental model, we injected LPS intracortically into Cx3CR1^+/GFP^ (microglia reporter) mice. Multi-label immunostainings confirmed that Cx3CR1-GFP microglia coexpressing P2Y12R and CD45 displayed cytoplasmic IL-1α and IL-1β production (Figure 1D). These findings collectively suggested that microglia are likely to be major mediators of central bumetanide actions under inflammatory conditions. Supporting this, central LPS administration resulted in a significant decrease of NKCC1 (encoded by *Slc12a2*) mRNA levels, along with a decline of P2Y12R mRNA, a key purinergic receptor reportedly associated with homeostatic microglial actions, microglial activation and response to injury (Figure 1E) [31,44–46].

### Deletion of microglial NKCC1 markedly impacts on baseline cell morphology and alters transformation to reactive microglia

Available RNA-seq data demonstrate that the *Slc12a2* gene is expressed by microglia [21, 40]. In line with this, our qPCR data revealed that mRNA levels of NKCC1 in microglial cells isolated from newborn (1.052±0.054) or adult (0.89±0.098) C57BL/6J mice are comparable to those in neural progenitors derived from embryonic brains (1.30±0.05; Figure 2A). However, NKCC1 mRNA expression markedly decreased when cells were grown in culture for 10 days (Figure 2A), making it difficult to study NKCC1 function in microglia using *in vitro* techniques.

**Figure 2.**
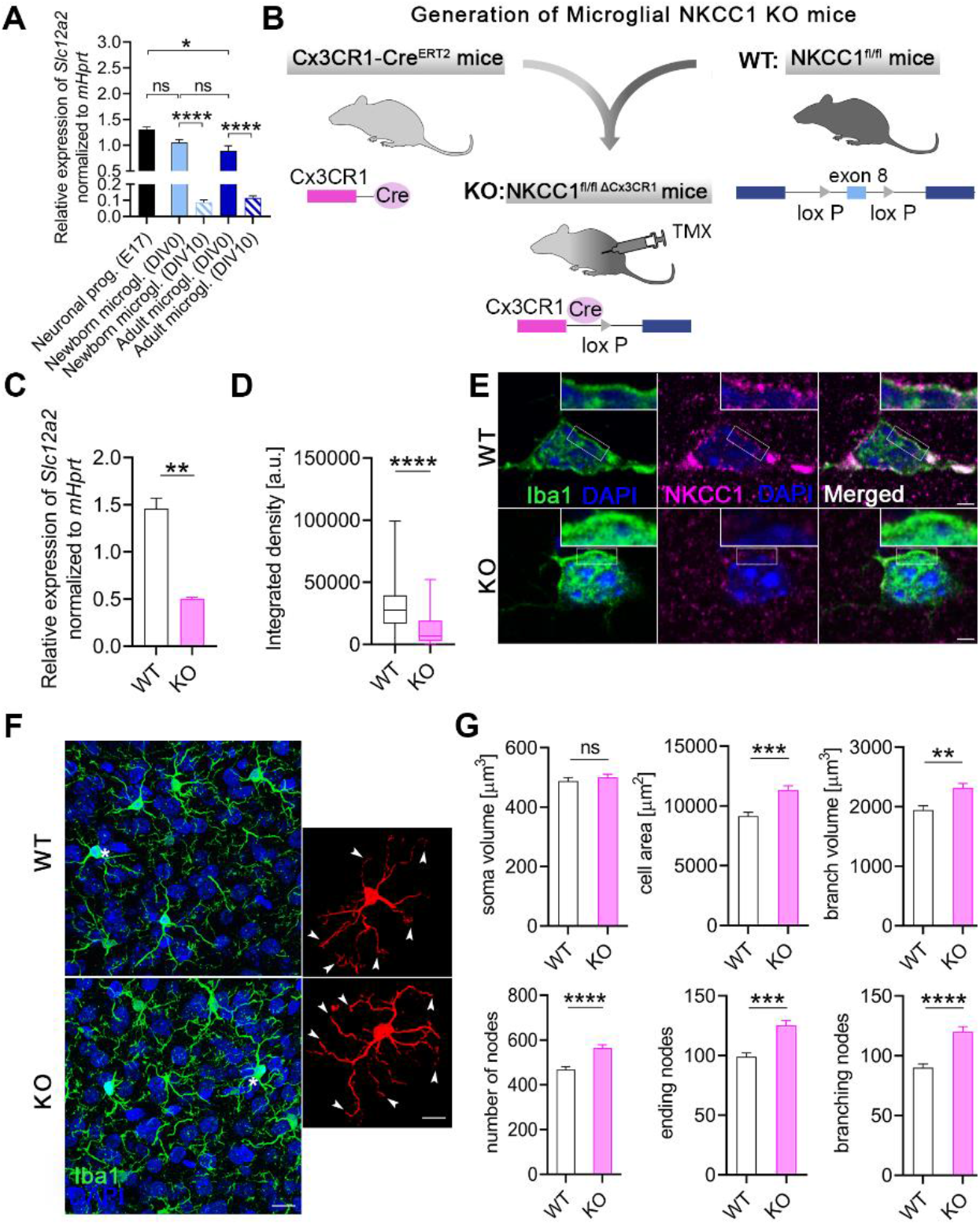
Deletion of microglial NKCC1 markedly impacts on baseline cell morphology and alters transformation to reactive microglia. **A:** NKCC1 mRNA expression levels in newborn and adult microglia isolated from C57BL/6J mice compared to neural progenitors derived from E17 embryonic hippocampi. Note, that NKCC1 mRNA levels decrease dramatically during *in vitro* maintenance (DIV10). **B:** We generated a novel microglia-specific conditional NKCC1 KO transgenic mouse line by crossing NKCC1^fl/fl^ (exon 8 of the *Slc12a2* gene was flanked with lox P sites) and Cx3CR1-Cre^ERT2^ mice. **C:** NKCC1 mRNA levels in isolated NKCC1 KO microglia was markedly reduced in comparison to wild-type cells. **D-E:** NKCC1 protein expression in a large number of randomly sampled NKCC1 KO microglia cells is markedly reduced compared to WT cells. Inserts show plasma membrane localization of NKCC1. **F-G:** Automated morphological analysis and maximum intensity projections of confocal images. Inserts show cells marked with white asterisks in 3D. Arrowheads indicate altered branch structure of NKCC1 KO microglia. Automated morphological analysis shows that features of NKCC1 deficient microglia significantly differ from WT microglia. **A:** One-way ANOVA, followed by Holm-Sidak’s post hoc test. N=3/group. **: *p*<0.01; n.s.: not significant **C:** Mann-Whitney test, N=3/group. **: *p* <0.01 **D:** Mann-Whitney test, N (WT)=142 cells from 2 mice, N (KO)=83 cells from 1 mouse. ****: *p*<0.0001 **E:** Scale: 2 μm **F-G:** Scale: 10 μm; Mann-Whitney test, N (WT)=78 cells from 3 mice, N (KO)=136 cells from 5 mice. **: *p* <0.01, ***: *p* <0.001, ****: *p* <0.0001 Abbreviations: DIV: days in vitro; n.s.: not significant; TMX: tamoxifen

To study the functional role of NKCC1 in microglia *in vivo*, we generated a novel transgenic mouse line in which microglial NKCC1 was deleted (Figure 2B). *Slc12a2* expression in isolated microglia derived from tamoxifen-inducible microglial NKCC1 KO mice was markedly reduced in comparison with wild type littermates (WT: 1.46±0.11, KO: 0.50±0.016; Figure 2C) as assessed by qPCR. Reduced NKCC1 protein expression was also confirmed by unbiased fluorescent density measurements after systematic random sampling of a great number of individual microglial cells from Iba1/NKCC1 stained slices. In fact, a discrete NKCC1-labeling was seen in the plasma membrane of microglial cells in WT mice, which was absent in microglial NKCC1 KO mice (Figure 2D-E).

Microglia are known to respond to physiological and pathological challenges with fast morphological transformation [47], while the role of NKCC1 in cell volume regulation has been highlighted in previous studies concerning other cell types [48, 49]. To study the role of NKCC1 in microglial cell shape, we performed automated morphological analysis using perfusion-fixed brain sections obtained from WT and microglial NKCC1 KO mice (Figure 2F-G). First, we aimed to test whether microglial NKCC1 deficiency has any impact on baseline cell morphology. Our data revealed that NKCC1-deficient microglia displayed a 23.8% higher cell surface, 19% higher branch volume, and 33.5% more branches compared to WT cells (Figure 2F-G), while we observed no changes in the cell body volume of KO versus WT microglia (Figure 2G). According to these data, microglial NKCC1 is likely to be involved in shaping baseline cell morphology.

### Microglial process motility is altered by central NKCC1 inhibition

To assess whether NKCC1 is involved in the regulation of dynamic microglial process surveillance and injury-induced microglial responses, we performed *in vivo* two-photon imaging in microglia reporter Cx3CR1^+/GFP^ mice in combination with bumetanide treatment (Figure 3). We tracked pre-lesion (baseline) process motility followed by focal laser-induced lesioning, which was repeated after bumetanide administration into the *cisterna magna* in a different (undisturbed) part of the cerebral cortex in the same animals (Figure 3A-B). Bumetanide caused a small, but significant (7%) increase in the mean velocity of microglial processes (Figure 3C). In contrast, however, lesion-induced recruitment of microglial processes showed a marked, 26,6% reduction after bumetanide treatment (N=6) (Figure 3D-E and see **Supplementary Video 1**). These experiments indicate that beyond shaping cell morphology, microglial NKCC1 also regulates dynamic microglial actions both under physiological and pathological conditions.

**Figure 3.**
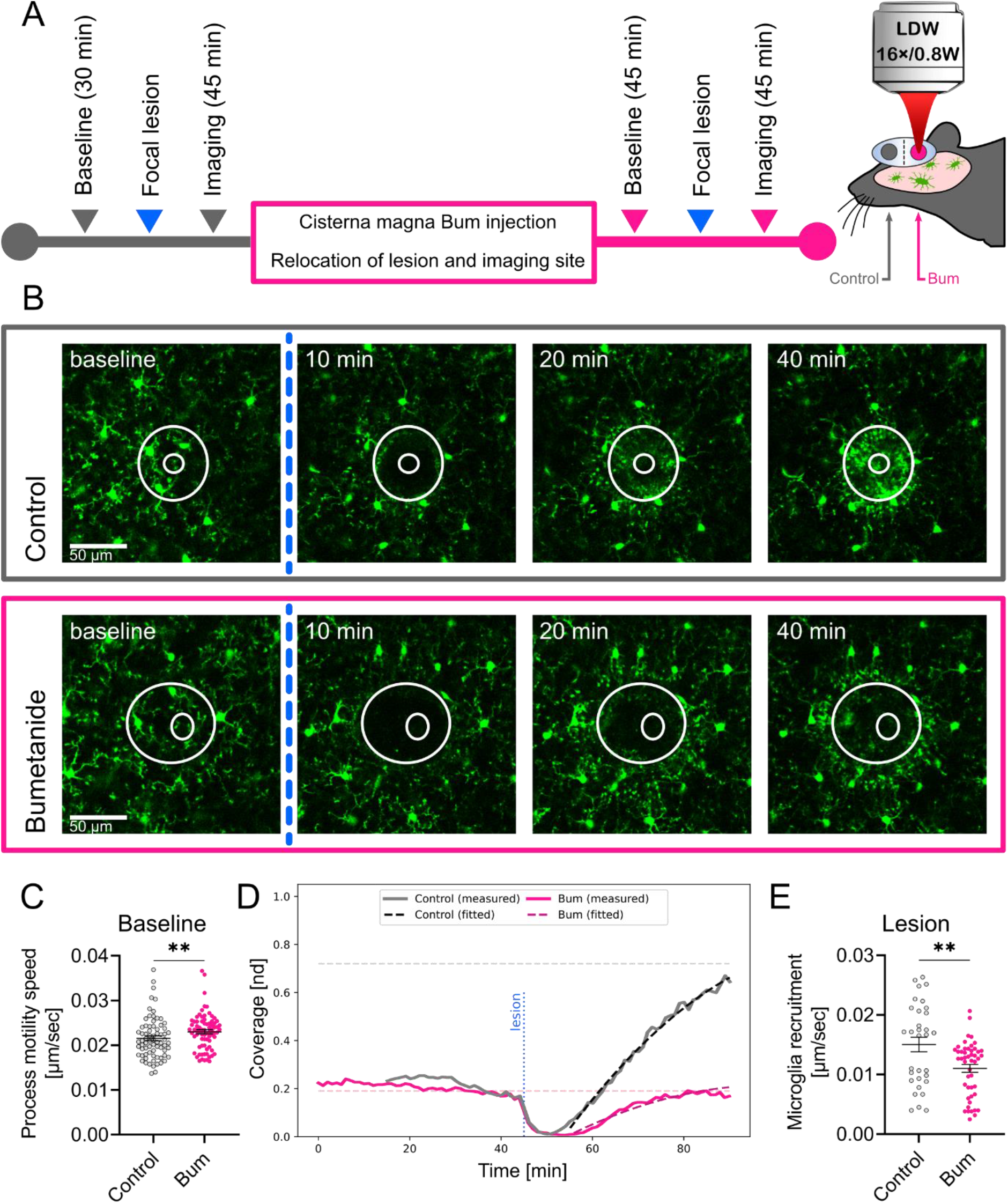
Microglial process dynamics and injury-induced process recruitment are altered by bumetanide. **A**: Outline of the 2-photon imaging experiment performed in Cx3CR1^+/GFP^ (microglia reporter) mice. Imaging was performed to assess baseline microglial process motility and response to laser-induced injury followed by bumetanide administration into the cisterna magna and identical measurements at a different, undisturbed cortical region. **B:** Representative images of microglial responses at selected time points, taken from Supplementary Video 1. The lesion site is marked with circles, but the inner zone of the lesion was excluded from analysis (see Materials and Methods). **C:** Mean base-state velocity distribution of processes from manual image tracking. Data show a 7% higher mean velocity of cell processes after Bum administration. **D:** Evaluation of microglial process coverage over time in the lesion-centered circular area generated by a custom image analysis pipeline. Solid lines show the proportion of the area covered by microglial GFP signal. Dashed lines show the calculated values of coverage from the best-fitting curves. Horizontal lines are the predicted maximal coverage values for the lesion site, vertical line is the time point of the lesioning. **E:** Calculated post-lesion velocities of microglial process recruitment using automated image analysis followed by model fitting. Data show a significant decrease (26,6%) in mean frontline velocity of cells with Bum administration. **C:** Mann-Whitney test, N_Control_=73, N_Bum_=76 processes from 5 mice, **: *p* <0.01 **E:** Mann-Whitney test, N_Control_=32, N_Bum_=48 fitted values from 4 and 6 mice, respectively, **: *p* <0.01; Bum: bumetanide

### The absence of microglial NKCC1 induces NLRP3 and potentiates inflammatory cytokine production in the cerebral cortex

To assess the effect of NKCC1 deletion on microglial inflammatory states and responses, we first examined the expression of IL-1β and NLRP3. Assembly of the NLRP3 inflammasome is known to be a key step for processing of pro-IL-1β by caspase-1, and the release of mature IL-1β from microglia or macrophages [50, 51]. qPCR data revealed elevated expression of both IL-1β and NLRP3 in NKCC1 KO microglia even in the absence of inflammatory stimulus (Figure 4A). Next, we examined the impact of NKCC1 deficiency on morphological characteristics of microglia after activation by endotoxin that primes microglial inflammatory responses and leads to morphological transformation [52]. Unbiased, automated morphological analysis showed that both WT and NKCC1 KO microglia respond to intracortical LPS administration with dramatic morphological conversion (Figure 4C). However, NKCC1-deficient microglia were significantly smaller than their wild type counterparts, displaying a 10% smaller cell surface, 12.5% smaller cell volume and 13.7% smaller branch volume (Figure 4D). According to these data, microglial NKCC1 is likely to be involved in shaping cell morphology and regulation of cell volume after inflammatory challenges.

**Figure 4.**
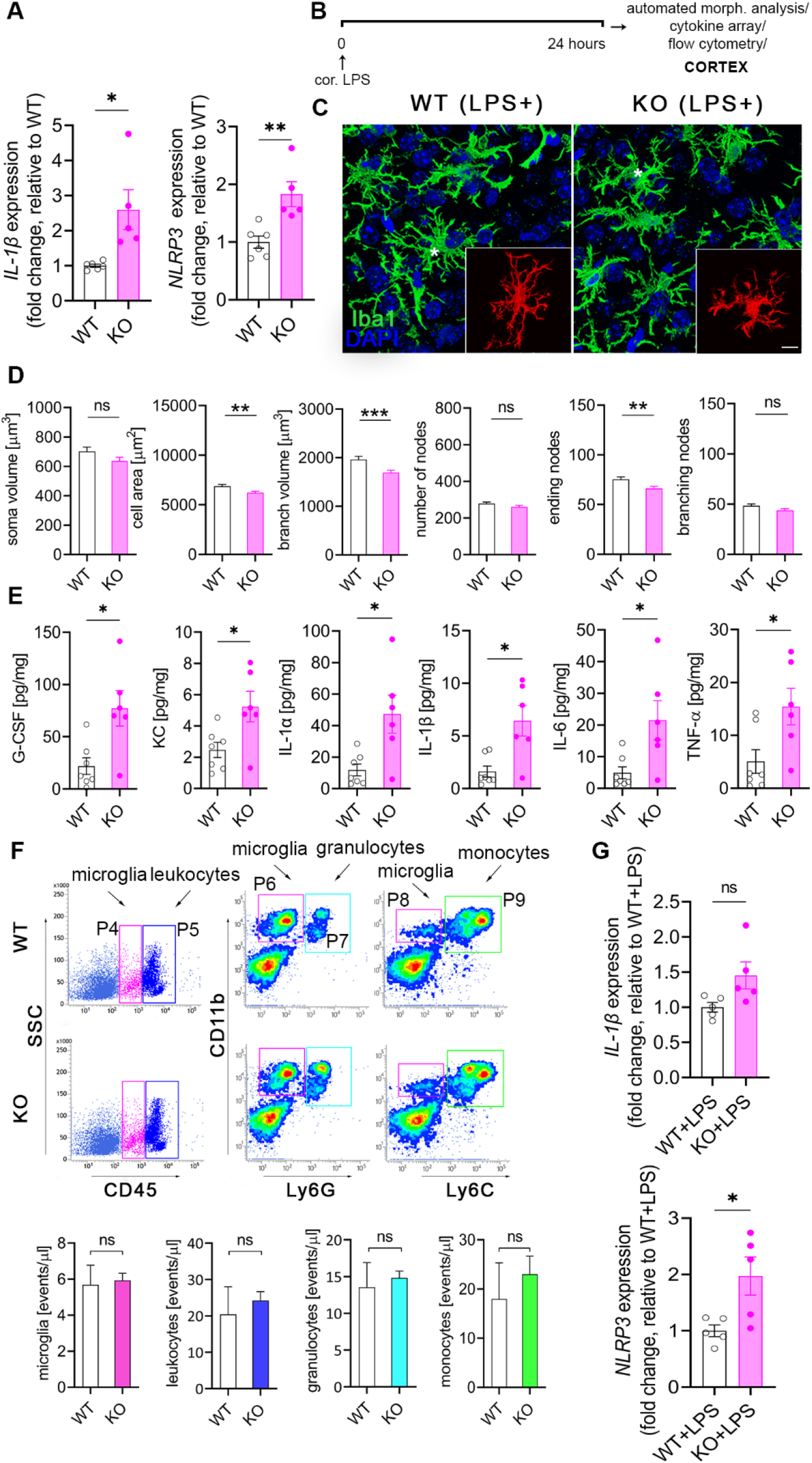
The absence of microglial NKCC1 boosts inflammatory cytokine production in the cerebral cortex in response to an inflammatory stimulus. **A**: Baseline NLRP3 and IL-1β mRNA expression is increased in isolated NKCC1 KO microglia compared to WT cells. **B:** Experimental outline of automated morphological analysis, cytokine array and flow cytometry. **C-D**: Automated morphological analysis show that activated NKCC1 deficient microglia are slightly smaller than their wild-type counterparts. **E**: LPS-induced cytokine levels are significantly higher in the cortices of microglial NKCC1 KO mice than in WT. **F:** Flow cytometric dot plots show that microglial NKCC1 deficiency does not alter the number of CD11b^+^, CD45^int^ microglia (P4 gate) or numbers of infiltrating CD11b^+^, CD45^high^ leukocytes (P5 gate), CD11b^+^, Ly6C^high^ monocytes (P9 gate) and CD11b^+^, Ly6G^high^ granulocytes (P7 gate) in response to intracortical LPS administration. See corresponding data on peripheral cytokine levels and immune cell populations in Supplementary Figure 3**. G:** Increased NLRP3 and IL-1β mRNA levels are sustained in NKCC1 KO and WT microglia 24 hours after intracisternal LPS administration**. A:** Unpaired t-test; \**p*<0.05, \*\**p*<0.01; N (WT)=6, N (KO)=5 **D:** Mann-Whitney test, N (WT)=171 cells from 6 mice, N (KO)= 85 cells from 4 mice, \*\**p*<0.01, \*\*\**p*<0.001 **E:** Mann-Whitney test, *: *p* <0.05; N (WT)=7, N (KO)=6 **F:** unpaired t-test, N (WT)=4, N (KO)=4 **G:** Unpaired t-test; \**p*<0.05; N (WT)=5, N (KO)=5; n. s.: not significant.

To test the impact of microglial NKCC1 deletion on the production of central inflammatory mediators, we examined LPS-induced cytokines and chemokines in microglial NKCC1 KO and WT mice twenty-four hours after intracortical LPS injections in brain homogenates (Figure 4E). Importantly, deletion of microglial NKCC1 markedly potentiated LPS-induced levels of G-CSF (3.53-fold), KC (2.12-fold), IL-1α (3.98-fold), IL-1β (3.96-fold), IL-6 (4.35-fold) and TNF-α (3-fold) in the cerebral cortex compared to controls (Figure 4E). In contrast, intracortical LPS treatment did not alter cytokine levels or T cell, B cell, monocyte or granulocyte numbers in the spleen or liver (Supplementary Figure 3A-C). Similarly to that seen after intracortical pharmacological inhibition of NKCC1 by bumetanide, neither the number of CD11b^+^ CD45^int^ microglial cells nor blood-borne CD11b^+^ CD45^high^ leukocytes, CD11b^+^ Ly6C^high^ monocytes or CD11b^+^ Ly6G^high^ granulocytes were altered as a result of microglial NKCC1 deficiency (Figure 4F). We also tested whether increased baseline IL-1β and NLRP3 mRNA levels seen in NKCC1 KO microglia (Figure 4A) were maintained in these cells 24 hours after intracisternal LPS administration. We found that significantly increased NLRP3 mRNA levels (by 100%) were still present in NKCC1 KO microglia isolated from the brain by using magnetic separation, while a strong trend (p=0.057) to increased IL-1β mRNA by 50% was also observed (Figure 4G). These findings corroborated our previous results on cortical NKCC1 blockade by bumetanide, confirming that the inhibition of microglial NKCC1 markedly potentiates central cytokine production upon inflammatory challenges, with a likely role for the NLRP3 inflammasome in the excessive production of IL-1β.

### Deletion of NKCC1 from microglia results in altered membrane currents and decreased intracellular Cl^-^ concentration

To test whether NKCC1 deletion changes ionic gradients across the microglial plasma membrane and alters intracellular ion concentrations, perforated patch-clamp recordings were performed on microglial cells in acute hippocampal slice preparations from WT and NKCC1 KO animals (Figure 5A). This method allows the measurement of membrane currents while intracellular Cl^-^ concentration remains unchanged. To measure membrane currents, recordings were carried out in voltage-clamp mode at -40 mV holding potential, while a pulse train of voltage steps (−100 mV to 100 mV with 20 mV increments and 100 ms duration) were delivered to construct I-V plots for individual microglial cells. (Figure 5B). I-V curves were obtained both in normotonic and in hypotonic ACSF solution (50% dilution, recorded after 5 min of perfusion). As previously described, an osmotic change due to a hypotonic medium causes microglial cells to stretch, activating volume-regulated anion channels (VRACs) and inducing an outwardly-rectifying current that is mainly driven by Cl^-^ ions [53–55]. Our measurements showed that both in normotonic and in hypotonic conditions, membrane currents evoked by voltage steps are markedly reduced in microglia with deletion of NKCC1, compared to ones in WT (Figure 5C; WT: N=8, KO: N=8 cells). We also found that resting membrane potentials of microglia in the normotonic condition are significantly more hyperpolarized in the NKCC1 KO compared to WT (Figure 5D; WT: N=13, KO: N=12 cells). Furthermore, reversal potential of swelling-induced current (calculated by subtracting the normotonic I-V curve from the hypotonic one) tend to be significantly lower in KO than in WT (Figure 5E). Estimations of intracellular Cl^-^ ion concentrations based on the swelling-induced current (calculated using the Nernst equation and the reversal potential values of the currents) also suggest that Cl^-^ ion concentration in microglia lacking NKCC1 is lower than in WT. (Figure 5F).

**Figure 5.**
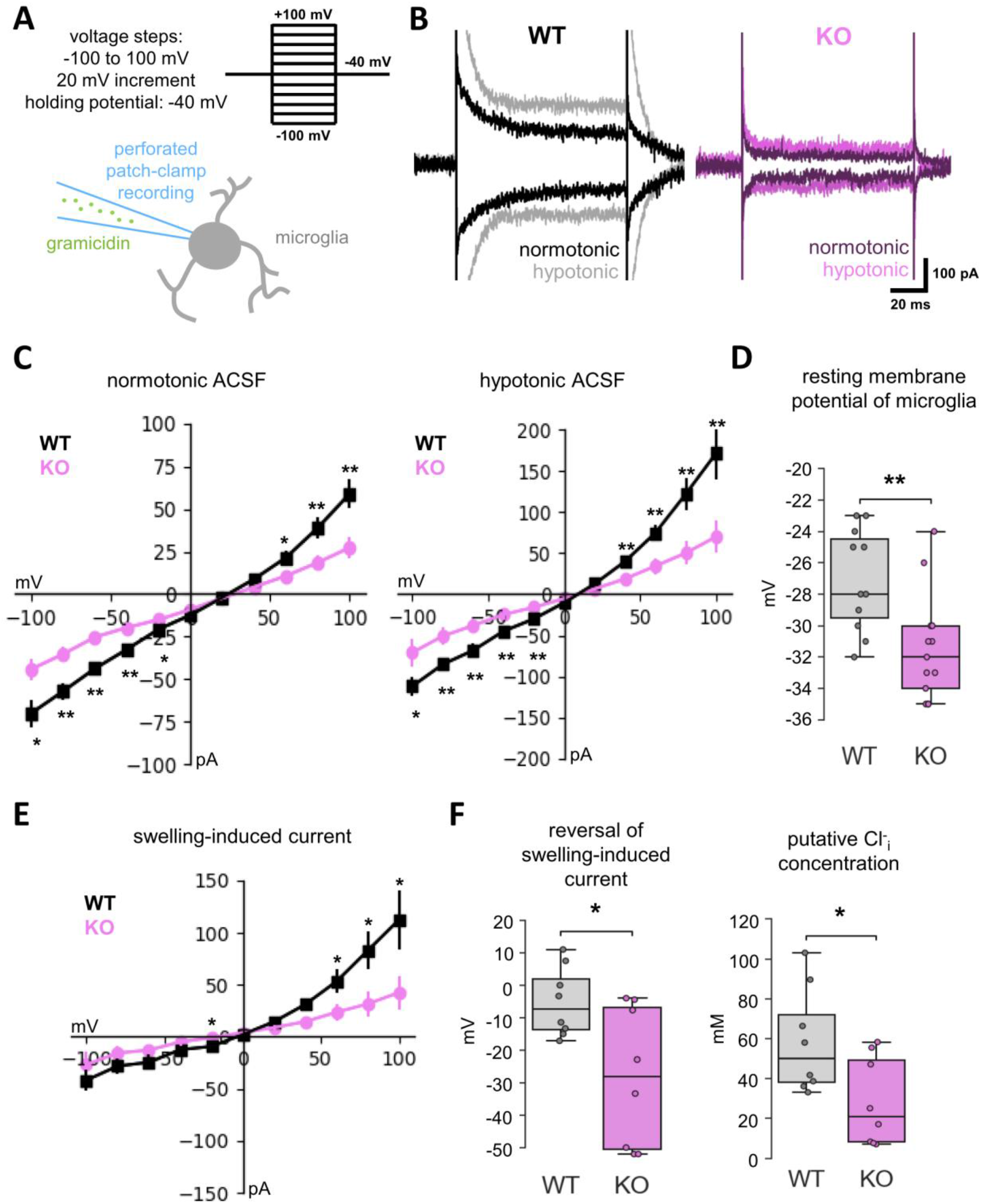
Deletion of NKCC1 from microglia results in altered membrane currents and decreased intracellular Cl- concentration. **A**: Schematic representation of experiment. Perforated patch-clamp recordings were performed on microglial cells in acute hippocampal slice preparations. Current responses to a train of voltage steps from -100 to 100 mV with 20 mV increments and a duration of 100 ms were measured in voltage-clamp mode (holding potential: -40 mV) both in normotonic, and after 5 minute perfusion with hypotonic ACSF (50% dilution). **B:** Example traces of recordings from WT (black: normotonic, grey: hypotonic ACSF) and NKCC1 KO (purple: normotonic, violet: hypotonic ACSF) animal. Traces show responses for -100 and +100 mV stimulations in both conditions. **C:** Average I-V curve responses from WT (black squares with SEM) and NKCC1 KO cells (violet circles with SEM) in normotonic (left) and after 5 minute perfusion of hypotonic ACSF (right) **D:** Resting membrane potential in normotonic condition of WT vs. NKCC1 KO microglial cells. **E:** I-V curves calculated by the subtraction of measured values in normotonic conditions from ones in hypotonic medium, resulting in I-V curves representing the currents evoked by cell-swelling due to osmotic change. **F:** Reversal potentials of the swelling-induced currents measured from WT (grey) or NKCC1 KO (violet) animals (left), with corresponding intracellular Cl^-^ concentrations (right) calculated via the Nernst equation. All parametric data are expressed as mean±SEM. **C:** Unpaired t-test; N(WT)=8 cells from 7 animal, N(KO)=8 cell from 6 animal; *: p<0.05, **: p<0.01 **D:** Unpaired t-test; N(WT)=13 microglial cells from 12 mice, N(KO)=12 microglial cells from 11 mice; **: *p*<0.01 **E:** Unpaired t-test; *: *p*<0.05 **F:** Unpaired t-test; N(WT)=8 cell, N (KO)=8 cell; *: p<0.05.

To investigate the potential compensatory mechanisms that may arise in response to NKCC1 deletion in microglial cells, we evaluated the expression of some relevant genes that are reportedly expressed by microglia and are known to regulate ion concentrations, membrane potential or response to osmotic stress (Supplementary Figure 4.) *Slc9a1, Slc8a1, Slc12a6, Clic1, Clcn3, Kcnk13, Kcnk6, Kcnj2, Kcna3, and Sgk1* mRNAs did not show altered expression between WT and NKCC1 KO microglia, but we found a more than two-fold increase in Lrrc8d, the D subunit of volume regulated anion channel (VRAC) in response to NKCC1 deletion (Supplementary Figure 4A). Importantly, Lrrc8d (VRAC) is involved in increasing the permeability for a broad range of uncharged organic osmolytes [56]. Interestingly, intracisternal LPS treatment resulted in a marked reduction of NKCC1 and P2Y12R mRNA levels (Figure 1E), in line with the downregulation of *Slc9a1*, *Slc8a1*, *Lrrc8d*, *Clic1*, *Clcn3*, *Kcnk13*, *Kcnk6*, *Kcnj2*, *Sgk1* genes (Supplementary Figure 4B) of which *Slc9a1, Clic1, Kcna3, Kcnj2, Kcnk13* genes are known to play a role in maintaining ramified morphology and resting state under physiological conditions [35,39,57,58].

### Deletion of microglial NKCC1 increases infarct volume, brain edema and worsens neurological outcome after MCAo

Because Na^+^-coupled Cl^-^ importers and their upstream regulatory serine-threonine kinases (WNK-SPAK-OSR1) are involved in maintaining intracellular ionic homeostasis as well as regulation of cell volume [2, 49], inhibiting these co-transporters is a subject of interest in ischemic stroke therapy and in other forms of acute brain injury (brain trauma, SAH) where cerebral edema is a major contributor to poor clinical outcome [2, 49]. Thus, we investigated whether microglial NKCC1 deficiency influences the severity of brain injury after experimental stroke in microglial NKCC1 KO mice subjected to transient, 45 min long middle cerebral artery occlusion (MCAo). Twenty-four hours after reperfusion, elimination of microglial NKCC1 was not associated with obvious alterations in astroglial GFAP levels or AQP4 levels in perivascular astrocyte endfeet (Supplementary Figure 5). However, we observed a significant increase in brain edema, which was accompanied by a 47% increase in lesion volume (Figure 6A-B; *p*=0.0262) and poor neurological outcome in NKCC1 KO mice (Figure 6B; Bederson’s score: WT: 1.17±0.17, KO: 2.0±0.24; Composit neurological score: WT: 14.9±1.65, KO: 21.9±2.12). Furthermore, experimental stroke induced a 4-fold upregulation of IL-1α and 4.8-fold increase in IL-1β expressing NKCC1 KO microglia cells compared to WT animals, while the overall microglia density and the number of activated microglia were not different at 24 hours reperfusion (Figure 6D-E). We also assessed whether increases in IL-1 production take place early after brain injury before the majority of the infarct is formed. Cytometric bead array did not reveal altered cytokine production in cortical homogenates shortly (8 hours) after MCAo (Figure 6C). Taken together, these results indicate that microglial NKCC1 deficiency results in augmented brain inflammation, brain edema and increased brain injury after experimental stroke.

**Figure 6.**
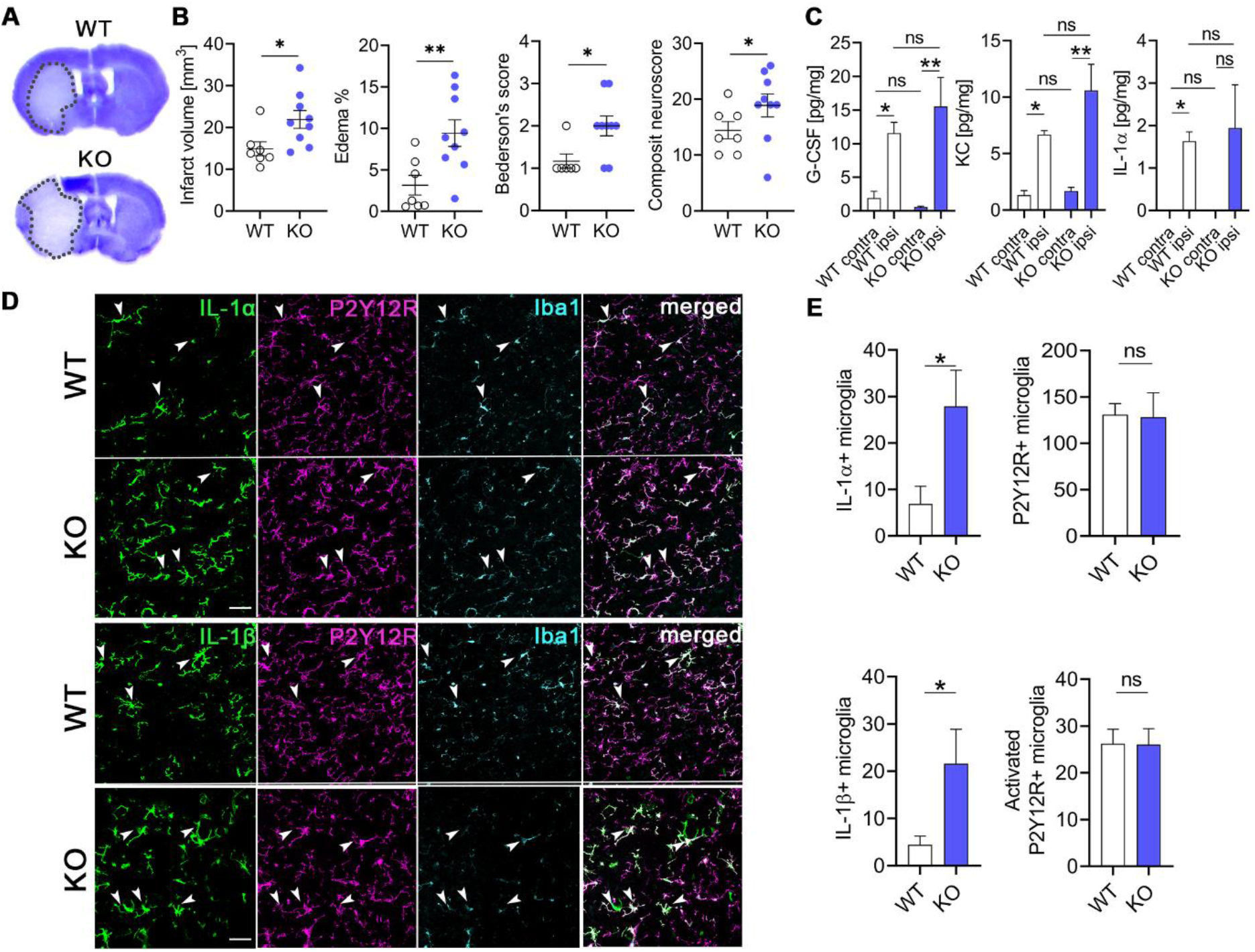
Deletion of microglial NKCC1 increases infarct volume, brain edema and worsens neurological outcome after MCAo. **A-B**: Microglial NKCC1-deficient mice (KO) show larger infarct volume as assessed on cresyl violet-stained brain sections and more severe neurological outcome compared to wild type mice. **C:** Cytokine levels in the cortex do not differ 8 hours after MCAo in KO mice compared to WT. **D-E:** Microglial NKCC1 deletion results in higher levels of IL-1α and IL-1β 24 hours after MCAo. **B:** Unpaired t-test, N (WT)=7, N (KO)=9; *: *p*<0.05, **: *p*<0.01 **C:** Kruskall-Wallis test followed by Dunn’s multiple comparison test; N=6/group. *: *p*<0.05, **: *p*<0.01 **D:** Scale: 50 μm **E:** Mann-Whitney test, *: *p*<0.05; N (WT)=7, N (KO)=9. n.s.: not significant

## Discussion

To our knowledge, we show for the first time that NKCC1 is functionally active in microglia in the adult mouse brain and that microglial NKCC1 is involved in shaping both baseline and reactive microglia morphology and responses. We present evidence that NKCC1 regulates microglial membrane potential, chloride levels and adaptation to osmotic stress in a cell-autonomous manner, while the lack of NKCC1 results in primed microglial inflammatory states as evidenced by elevated NLRP3 and IL-1β levels. In line with this, while systemic NKCC1 blockade attenuates LPS-induced inflammation in the brain, central pharmacological inhibition or genetic deletion of microglial NKCC1 potentiates inflammatory cytokine production. Microglial NKCC1 deletion also increases brain injury, inflammation, cerebral edema, and leads to worse neurological outcome after acute brain injury induced by experimental stroke. Thus, microglial NKCC1 emerges as an important regulator of central inflammatory responses.

As pointed out in the Introduction, there is little direct information on whether systemically-applied bumetanide might exert therapeutic effects by directly blocking NKCC1 in the brain and, particularly in central neurons. To unravel the effects of pharmacological inhibition of NKCC1 in the periphery and the brain, we investigated the systemic vs. central effects of bumetanide in response to bacterial endotoxin (LPS) administration, which is widely used to trigger inflammatory cytokine responses [41, 52]. Intraperitoneally injected bumetanide reduced intracortical LPS-induced proinflammatory cytokine/chemokine responses in the brain, similarly to that found in the case of LPS-induced lung injury attributed to NKCC1-mediated macrophage activation [59]. These findings are also consistent with the suggested beneficial therapeutic effects of systemic NKCC1 blockade in CNS pathologies, which are associated with inflammation [1,7,8,10–12,60].

Importantly, intracortical administration of bumetanide had exactly the opposite effect on intracortical LPS-induced cytokine production to that seen after systemic bumetanide treatment, resulting in elevated cytokine levels in the cortex. These markedly different outcomes clearly demonstrate the difficulty to interpret pharmacological interventions in the absence of data on the precise cellular targets of NKCC1 actions in the brain and highlight the need for cell-specific NKCC1 targeting studies.

Because microglia are key source of inflammatory cytokine in the brain, we tested whether microglial activity could be directly related to NKCC1 function. Indeed, we observed comparable *Slc12a2* expression in microglia, isolated from either newborn or adult mouse brains, to that seen in neural progenitors. We also revealed NKCC1 protein in cortical microglial cells, which is in line with recent observations showing microglial NKCC1 expression in the superficial spinal dorsal horn in rats [61], as also supported by single-cell transcriptomic studies [21, 40]. Further on, we demonstrated that NKCC1 levels are markedly downregulated in microglia in response to LPS, alongside with decreased P2Y12R levels (a purinergic receptor regulating microglial cell-cell interactions in response to injury [21,44,62– 64]), suggesting that impaired microglial NKCC1 function alters microglial responses to injury or inflammatory stimuli.

To investigate the cell-autonomous effects of NKCC1 in microglia, we generated a new, microglial NKCC1 KO transgenic mouse line. We found that the absence of NKCC1 resulted in a more branching microglial morphology in resting state and a more amoeboid shape under inflammatory conditions. These results paralleled enhanced baseline process motility in the intact brain, *in vivo,* and significantly reduced process recruitment to the site of the acute lesion, when microglial NKCC1 was blocked by intracisternal bumetanide during two-photon imaging. Associations of NKCC1 with actin cytoskeleton dynamics [65, 66] are well-documented, which take place via controlling F-actin organization through Cofilin-1 and RhoGTPase activity [65]. Thus, our results suggest that regulation of targeted process outgrowth in microglia is NKCC1-dependent, which is known to depend on microglial P2Y12R, but does not require THIK-1, a potassium channel that regulates baseline microglia shape and process motility [35].

Importantly, we also found that the absence of microglial NKCC1 renders microglia to express higher levels of NLRP3 and IL-1β mRNA even without any exposure to inflammatory stimuli. It is well-established that changes in microglial membrane potential and ion currents markedly determine cell volume regulation, process dynamics and inflammatory responses of microglia. For example, depolarization is associated with reduced branching in microglia [35, 67], while potassium or chloride efflux are essential for NEK7-NLRP3 interaction during the assembly of the NLRP3 inflammasome [38, 50], which is required for the production of mature IL-1β [37,38,50]. The regulation of these processes appears to be highly complex. The TWIK2 two-pore domain K^+^ channel (K_2P_) mediates K^+^ efflux triggering NLRP3 activation in lung macrophages [37], while THIK-1, has been identified as the main K^+^ channel in microglia [35], although RNA profiling [21, 40] also revealed high expression of TWIK2 in this cell type. In line with this, blocking THIK-1 function inhibits the release of IL-1β from activated microglia, consistent with potassium efflux being necessary for inflammasome activation [35].

To investigate how microglial NKCC1 may contribute to these processes, we first investigated whether deletion of NKCC1 leads to altered expression of ion channels or transporters that have been linked with microglial ion- and volume regulation and with inflammatory cytokine production (see references in Supplementary Figure 4D). We could not observe changes in THIK-1 or TWIK2 mRNA levels by qPCR, but the subunit D of volume regulated anion channel (VRAC) was upregulated in KO animals. Subunit D is involved in the transport of uncharged organic osmolytes by VRAC and it may be important for volume-reduction without a major release of inorganic ions [56] for a cell coping with potentially lowered inorganic ion levels due to NKCC1 loss. Interestingly, LRRC8A-containing anion channels are also known associate with NADPH oxidase 1 (Nox1) and regulate superoxide production and TNF-α signalling in vascular smooth muscle cells [56, 68]. Thus, changes in microglial ROS production in the absence of NKCC1 will need to be investigated in future studies.

Next, we conducted perforated patch-clamp recordings on microglia in acute hippocampal slice preparations from both WT and KO animals. Importantly, we found lower intracellular Cl^-^ concentrations in the case of NKCC1 KO with resting membrane potentials significantly more hyperpolarized, while membrane currents evoked by voltage steps were markedly reduced in NKCC1 KO microglia under both normotonic and hypotonic conditions. Thus, morphological differences in NKCC1 KO microglia may be a direct consequence of altered trafficking of Na^+^, K^+^ and Cl^-^ ions through the microglial membrane, which could lead to a more hyperpolarized membrane potential, more ramified morphology, and interference with cell volume regulation and inflammasome activation. Concerning the latter, we found that in line with increased NLRP3 and IL-1β mRNA levels in NKCC1 KO microglia, LPS stimulation resulted in elevated NLRP3 mRNA levels even after 24h upon LPS treatment, and potentiated the production of IL-1β and other cytokines in the cortex of microglial NKCC1 KO mice compared to control mice. Importantly, deletion of microglial NKCC1 had an identical effect on inflammatory cytokine levels to that seen after NKCC1 inhibition by bumetanide, suggesting that central bumetanide actions involve blockade of microglial NKCC1. In line with this, we could not find significant changes in two key astroglial markers, GFAP and AQP4 after central bumetanide application in response to LPS treatment, arguing against a major role of astroglial NKCC1 in the observed differences on the timescale of the present study. While a role for astroglial NKCC1 in mediating central bumetanide actions has been previously reported [18, 19], our present work shows that central bumetanide effects involve a substantial microglial component via NKCC1.

Proinflammatory cytokine production, particularly the effects mediated by IL-1, are known to influence acute brain injury induced by brain trauma or stroke [69–71]. We found significantly increased infarct volume (by 47%) in microglial NKCC1 KO mice after experimental stroke comparable to a striking, 60% increase in infarct size caused by highly efficient, selective microglia elimination [32]. Microglia depletion resulted in dysregulated neuronal network activity, an effect that is likely to be mediated via elimination of protective microglia-neuron interactions at somatic purinergic junctions [31]. It remains to be investigated in future studies, whether microglia-neuron interactions are altered in microglial NKCC1 KO mice, which could shape neuronal injury via diverse mechanisms related to changes in neuronal excitability [29,30,32,33]. In line with this, the absence of microglial NKCC1 may have contributed to increased neuronal injury via increased IL-1 production, as suggested by the established pathophysiological role of IL-1-mediated actions in different forms of brain injury [69, 72].

In our experimental stroke studies, we could not detect increased cytokine production in the absence of microglial NKCC1 early (8h) after MCAo, while both IL-1α and IL-1β expression were markedly upregulated in NKCC1 KO microglia 24 hours after reperfusion. Following stroke, impaired cell volume regulation results in cytotoxic cell swelling and edema formation, which peaks beyond the first 24 hours in both patients and experimental animals [73, 74]. Cerebral edema was shown to be associated with increased phosphorylation of the SPAK/OSR1 kinases playing a key role in NKCC1 activation in various neural cell types [2,48,49] and a SPAK kinase inhibitor, ZT-1a, attenuated cerebral edema after stroke [75]. In line with this, NKCC1 is involved in homeostatic cell volume regulation [48, 49] and bumetanide attenuates edema formation in response to ischaemic or traumatic brain injury [2]. In light of these findings, our data showing impaired volume regulation and exaggerated cytokine production in NKCC1 KO microglia have a potentially high pathophysiological and clinical relevance.

In conclusion, we show that microglial NKCC1 plays a previously unrecognized role in shaping microglial phenotype and inflammatory responses. Strikingly, bumetanide-induced enhancement of neuroinflammation was mimicked by conditional deletion of microglial NKCC1, suggesting that microglial NKCC1 is a significant target of NKCC1 blockers, given that brain tissue levels of such drugs are sufficiently high [10,13,14]. Our results implying microglial NKCC1 in inflammation, brain edema formation and responses to injury suggest that microglial NKCC1 actions should be considered when studying the effects of NKCC1 inhibitors in different CNS functions and pathologies. This is further emphasized by the broad clinical interest in pharmacological NKCC1 blockade. The present results also call for a re-evaluation of the pharmacological effects of bumetanide in brain diseases with a significant inflammatory component.

## Materials and Methods

### Experimental animals

Experiments were performed on adult male C57BL/6J (RRID: IMSR JAX:000664), NKCC1^fl/fl^ and NKCC1^fl/fl Δ Cx3CR1^ (microglial NKCC1 knockout), mice (all on C57BL/6J background). For experiments using LPS treatment P90-110, and for MCAo experiments P70-90 day old mice were used. NKCC1^fl/fl^ mice were kindly provided by Dr. Christian A. Hübner of University Hospital Jena of Friedrich Schiller University, Jena, Germany. Ubiquitous NKCC1 KO mice (P15-20, NKCC1 full knockout, NKCC1^-/-^), generated by crossing C57BL/6NTac-Gt(ROSA)26Sor^tm16(cre)Arte^ and NKCC1^fl/fl^ mice, were used as immunostaining controls and were not subjected to other experimental procedures. Mice were housed in a 12 hours dark/light cycle environment, under controlled temperature and humidity and *ad libitum* access to food and water. All experimental procedures were in accordance with the guidelines set by the European Communities Council Directive (86/609 EEC) and the Hungarian Act of Animal Care and Experimentation (1998; XXVIII, Sect. 243/1998), approved by the Animal Care and Experimentation Committee of the Institute of Experimental Medicine and the Government Office of Pest Country Department of Food Chain Safety, Veterinary Office, Plant Protection and Soil Conservation Budapest, Hungary under the number PE/EA/1021-7/2019 and Department of Food Chain Safety and Animal Health Directorate of Csongrád Country, Hungary. All experiments followed ARRIVE and IMPROVE guidelines [76, 77]. A code was allocated to each animal using GraphPad’s random number generator and was randomly assigned to different treatment groups. During all surgical procedures and functional tests the experimenter was blinded to the treatment.

### Generation and genotyping of the microglial NKCC1 knockout mouse line

Microglial NKCC1 deletion (NKCC1^fl/fl ΔCx3CR1^) was achieved by crossing tamoxifen-inducible B6.129P2(C)-CX3CR1tm2.1 (cre/ERT2)Jung/J mice (RRID:IMSR_JAX:020940JAX) [78] with the NKCC1^fl/fl^ mouse line [79], in which a region between exon 7 and exon 10 of *Slc12a2* gene was flanked with lox P sites. Microglial NKCC1 deletion was achieved by two intraperitoneally administered tamoxifen injections (100 ul of 20mg/ml in corn oil, #T5648, Sigma-Aldrich) 48 hours apart in 3-4 weeks old male mice, shortly after weaning. In all functional experiments mice were studied more than 3 weeks after the last tamoxifen injection to avoid interference with Cre-dependent recombination in peripheral myeloid cells [78]. In the case of selective microglia isolation by magnetic cell sorting, we used animals exposed to tamoxifen for at least two weeks as Cre activity was shown to increase most dramatically within the first week after initiating treatment [80].

In all experiments, conditional mutant mice heterozygous for cre and homozygous for NKCC1 flox (^cre/+ fl/fl^), referred to as NKCC1 KO throughout the manuscript were used. In all cases, littermate controls - referred to as WT - were used which were negative for cre and homozygous for NKCC1 flox (^+/+ fl/fl^). Heterozygous, conditional mutant mice were viable, fertile, average in size, and did not display any gross physical or marked behavioral abnormalities. Genotypes for *Slc12a2* encoding NKCC1 were determined from tail biopsy samples by conventional PCR using the following primers: 5’- GCAATTAAGTTTGGAGGTTCCTT, 5’-CCAACAGTATGCAGACTCTC and 5’-CCAACAGTATGCAGACTCTC; product sizes: 200 bps for WT, 260 bps for floxed and 460 bps for KO.

### Microglia cell preparations and cultures

Microglia cells from either newborn (P0-1, male or female) or adult (P40-55, male) C57BL/6J, NKCC1^fl/fl^ (WT) or NKCC1^fl/fl Δ Cx3CR1^ (KO) mice were isolated by magnetic separation using anti-CD11b microbeads (Miltenyi Biotec, Germany), with slight modification of the protocol described by Otxoa-de-Amezaga [81]. After transcardial perfusion with ice-cold PBS (phosphate buffered saline), the brain tissues (cortices and hippocampi) were enzymatically dissociated with Neural Tissue Dissociation Kit-P (#130-092-628; Miltenyi Biotec). Myelin was removed by MACS Myelin Removal Beads II (#130-096-733, Miltenyi Biotec), then cells in a single-cell suspension were magnetically labeled with MACS CD11b microbeads (#130-093-634, Miltenyi Biotec) and were separated using MS columns (#130-042-201, Miltenyi Biotec, Germany). Cells selected with CD11b microbeads were plated onto poly-L-lysine precoated 96-well or 386-well plates at 3x10^4^ cell/cm^2^ density and were cultured at 37 °C in a 95% air/5% CO_2_ incubator in DMEM/Glutamax medium (#31966-021, Gibco) supplemented with 10% FBS (fetal bovine serum, #FB-1090, Biosera), 1% Pen/Strep (10000 U/ml; #15140-122, ThermoFisher Scientific) and 10 nM Macrophage Colony-Stimulating Factor (M-CSF; #PMC2044, ThermoFisher Scientific) for 10 days.

### Embryonic neural progenitor cell preparations

Embryonic hippocampal cells were prepared from C57BL/6J mice on embryonic day 17. After aseptically removing the hippocampi from the skull, tissue was freed from meninges and incubated in 0.05% trypsin-EDTA (#T4549, Sigma-Aldrich) solution with 0.05% Dnase I (#DN25, Sigma-Aldrich) in phosphate-buffered saline (PBS) for 15 min at 37°C.

### Intracortical and intracisternal injections

P90-110 mice were deeply anesthetized with fentanyl (0.05 mg/kg) and mounted into a stereotaxic frame, then were subjected to either single saline or LPS (lipopolysaccharide from *Escherichia coli* O26:B6; 200 nl of 5 mg/ml, rate=200 nl/10 min; #L8274, Sigma-Aldrich) injections using a glass micropipette [82]. The coordinates for the injection were anterior-posterior -2,5 mm, lateral +1,5 mm, and ventral -0,25 mm from the bregma. Bumetanide, a specific inhibitor of NKCC1 was co-injected with LPS (50 μM; #3108, Tocris). At 24h, mice were transcardially perfused with saline, and approximately 0.5x0.5x0.5 cm sized tissue pieces from the center of each injected cortical region were cut off and collected for cytokine array and flow cytometric analysis. For tissue sectioning mice were perfused with saline followed by 4% paraformaldehyde in PBS. For qRT-PCR experiments to assess the effect of NKCC1 deficiency on microglial expression of genes which contribute to ion homeostasis, membrane potential, cell volume regulation or inflammation, LPS (5 μg dissolved in ACSF) was administered into the cisterna magna in 5 μl final volume, using a glass capillary. At 24h, mice were transcardially perfused with saline followed by CD11b+ magnetic cell sorting.

### Systemic administration of LPS and bumetanide

Male adult NKCC1^fl/fl^ mice were injected intraperitoneally with saline, LPS (2 mg/kg; O26:B6, #L8274, Sigma-Aldrich) or LPS (2 mg/kg) and bumetanide (25 mg/kg; #3108, Tocris). Intraperitoneal bumetanide injections were repeated twice, the first one 15 min prior to LPS injection, the second one 1 hour after LPS administration. The double injection aimed to ensure that in the critical time window we have effective concentrations of bumetanide in the circulation. At 24h, saline-perfused spleen and brain samples were collected for cytokine measurements or flow cytometric analysis.

### Cytokine measurement

The levels of inflammatory cytokines and chemokines were measured in spleen and brain samples. Sample processing and protein determination were performed as described previously [83]. Mice were transcardially perfused with saline prior to the collection of spleen and brain samples (ipsilateral to injections). Tissue samples were homogenized in TritonX-100 and protease inhibitor-containing (1:100, #539131, Calbiochem) Tris-HCl buffer (TBS, pH 7.4) and centrifuged at 17000 g, for 20 min at 4 °C. Protein level was quantified for every sample using BCA Protein Assay Kit (#23225, ThermoFisher Scientific). Then, measured cytokine levels were normalized for total protein concentrations. The concentrations of cytokines and chemokines were determined by BD^TM^ Cytometric Bead Array (CBA) using BD CBA Flex Sets (GCSF: #560152, KC: #558340, IL-1α: #560157, IL-1β: #560232, IL-6: #558301, TNF-α: #558299, BD, Becton, Dickinson and Company) according to the manufacturer’s instructions. Samples were acquired using a BD FACSVerse flow cytometer (BD, Becton, Dickinson and Company) and the results were analyzed by FCAP Array software (BD, Becton, Dickinson and Company).

### Flow cytometric analysis of brain and spleen and liver samples

For flow cytometric analysis, cells were isolated from mouse brains with Collagenase D (0,5 mg/ml, #11088866001, Roche), DNase I (10μg/ml, #DN25, Sigma-Aldrich) dissolved in 10% FBS containing DMEM (#6546, Sigma-Aldrich), then the cell suspension was passed through a 40 μm cell strainer (Corning). After enzymatic dissociation, the cells were resuspended in 40% Percoll solution and overlayed on 70% Percoll (#17-0891-01, GE Healthcare). After a density centrifugation step (at 2100 rpm, 30 min), mononuclear cells were collected from the interphase of 40%/70% Percoll. Spleen was mechanically homogenized and red blood cells were removed by centrifugation. Before acquisition, brain, spleen and liver cells were diluted with FACS buffer and were incubated with anti-mouse CD16/32 to block Fc receptor. Brain cells or spleen and liver leukocytes were incubated with cocktails of selected antibodies: T cells - anti-mouse CD8a-PE (1:200, #12-0081-82, eBioscience), anti-mouse CD3-APC clone 17A2 (1:200, #17-0032-80, eBioScience), anti-mouse CD4-FITC (1:200, #11-0043-82, eBioscience), anti-mouse CD45-PerCP/Cy5.5 (1:200, #103131, BioLegend); B cells/granulocytes - anti-mouse CD19-FITC (1:200, #11-0193-81, eBioScience), anti-mouse Ly-6C-PE-Cy7 (1:500, #25-5932-80, eBioScience), anti-mouse Ly-6G-APC (1:500, #127613, BioLegend); monocytes/granulocytes - anti-mouse CD11b-FITC (1:200, #11-0112-81, eBioscience); anti-mouse Ly-6C-PE-Cy7; anti-mouse Ly-6G-APC, CD45-PerCP/Cy5.5. To exclude dead cells, some coctails contained propidium iodide (3 μM; #P1304MP, ThermoFisher). Cells were acquired on a BD FACSVerse flow cytometer and data were analyzed using FACSuite software (BD, Becton, Dickinson and Company). Cell counts were calculated by using 15 μm polystyrene microbeads (#18328-5, Polysciences).

### RNA isolation and cDNA reverse transcription

For qRT-PCR measurements, CD11b+ magnetic sorted microglia cells or embryonic neural progenitor cells were homogenized in QIAzol Lysis Reagent (#79306, QIAGEN) either immediately after isolation or after 10 days of in vitro culturing. Total RNA was isolated using Direct-zol™ RNA Miniprep Kits (#R2052, Zymo Research) following the manufacturer’s protocol. The RNA purity and concentration were assessed by NanoDrop ND-1000 spectrophotometer (Nanodrop Technologies). The isolated RNA was then stored at −80 °C. RNA was subjected to DNase I (#AM2224, Ambion) treatment in the presence of RNase H inhibitor (#AM2682, Ambion). Standardized quantities of RNA were reverse transcribed to cDNA using the SuperScript™ II First-strand Reverse Transcriptase system (#18064014, ThermoFisher Scientific) and random hexamers (#48190011, Invitrogen) supplemented with RNase H inhibitor (#AM2682, Ambion).

### Real-time quantitative PCR (qRT-PCR)

Real-time quantitative PCR was performed with QuantStudio12K Flex qPCR instrument (Applied Biosystems), using TaqMan Gene Expression Assays and the TaqMan Gene Expression Master Mix (#4369016, ThermoFischer Scientific). All mouse TaqMan Gene Expression Assays used for amplification reactions were obtained from ThermoFischer Scientific: *Slc12a2* (Mm01265951_m1, targeting exon 1-10); (Mm00436546_m1, targeting exon 8-10, used for the validation of microglia specific NKCC1 deletion, see Figure 2C), *Hprt* (Mm03024075_m1), *Slc12a6* (Mm01334052_m1), *Slc8a1* (Mm01232254_m1), *Slc9a1* (Mm00444270_m1), *Clcn3* (Mm01348786_m1), *Clic1* (Mm00446336_m1), *Kcnk6* (Mm01176312_g1), *Kcnj2* (Mm00434616_m1), *Kcna3* (Mm00434599_s1), *Lrrc8d* (Mm01207167_m1), *Sgk1* (Mm00441380_m1), *NLRP3* (Mm00840904_m1), *pro-IL-1β* (Mm00434228_m1). The amplification was performed under the following cycling conditions: 95 °C for 10 min, followed by 40 cycles of 95 °C for 10 sec, and 60 °C for 1 min. The comparative Ct method (ΔΔCt method) was used to analyze the relative expression values for each transcript using Hprt as a reference gene.

### In vivo two-photon imaging and assessment of microglial process dynamics

CX3CR1^+/GFP^ microglia reporter mice were anaesthetized using fentanyl. As previously has reported [31], fentanyl did not influence microglial process motility compared to the effects of different anaestethics. Cranial window with 3 mm diameter was opened on the left hemisphere centered 1.5 mm lateral and 1 mm posterior to bregma without hurting the dura mater. After removal of the skull bone a 3 mm and 5 mm double glass coverslip construct was fixed with 3M™ Vetbond™ tissue glue on top of the dura mater. Then a custom-made metal headpiece (Femtonics Ltd., Budapest, Hungary) was fixed with dental cement on the surface of the skull. All experiments were performed on a Femto2D-DualScanhead microscope (Femtonics Ltd., Budapest, Hungary) coupled with a Chameleon Discovery laser (Coherent, Santa Clara, USA). Following excitation, the fluorescent signal was collected using a Nikon 18X water immersion objective. Data acquisition was performed by MES software (Femtonics Ltd.). Galvano Z-stacks of 8 images (500x500 pixels, 3 μm step size, range=100-125 μm from pial surface) were made at every minute. Two-photon image sequences were exported from MES and analyzed using FIJI. Microglial baseline process velocity was measured on time-series images acquired with 2P microscopy. Following motion correction, images from the same region of CX3CR1^+/GFP^ mice were analyzed with the Manual Tracking plugin of FIJI. We applied a local maximum centring correction method with a search square of 5 pixels. Pixel size was 0.65 μm/px. Processes were included in the measurement when they were clearly traceable for at least 10 minutes. To compare how fast microglia cells are responding to injuries, cortical lesion was formed using laser beam. Focal lesion was induced with an 1040 nm fix laser.

Then, we created an automated image processing pipeline using CellProfiler [84] to determine the proportion of area over time covered by microglia cells on each image. The perimeter of the lesion in all image sets was marked manually, and only cells inside this region was taken into account, ignoring also some of the area in the middle as there were autoflourescent artifacts detected there. To differentiate between cells (high intensity areas) and background (low intensity areas), an adaptive threshold value was calculated from the per-image median intensities. The coverage was then measured as the proportion of pixels classified as cell and the total number of pixels in the image. Coverage values in the initial (baseline) phase of the experiment kept stable, no remarkable oscillation was recorded in any cases. Based on the observed data, we assumed that coverage reaches a minimum value 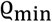 as a result of the injury, then at time *t_s_*, microglia cells start to flow into the lesion area uniformly from its perimeter with velocity *v* and saturate at a final coverage 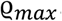. From this assumption, we created a simple mathematical model that predicts the coverage values over time inside the lesion area.

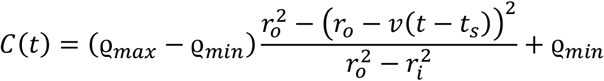

In this equation, *r_i_* and *r_o_* denote the inner and outer radii of the ring-shaped lesion site, respectively. See the inserted drawing for a visual explanation of the model. For each measured coverage dataset, we fitted this model by choosing values for *v*, *q_max_* and *t_s_* that minimize the mean squared difference between the observed and predicted values. To find the best fitting values, a grid search-based optimizer algorithm was used. We found that even this simple model is able to accurately describe the observed behavior of cell flow from the external areas inside the lesion, and it also gives an estimate for the cell’s mean velocity which is in good agreement with those from the manual tracking of cell processes.

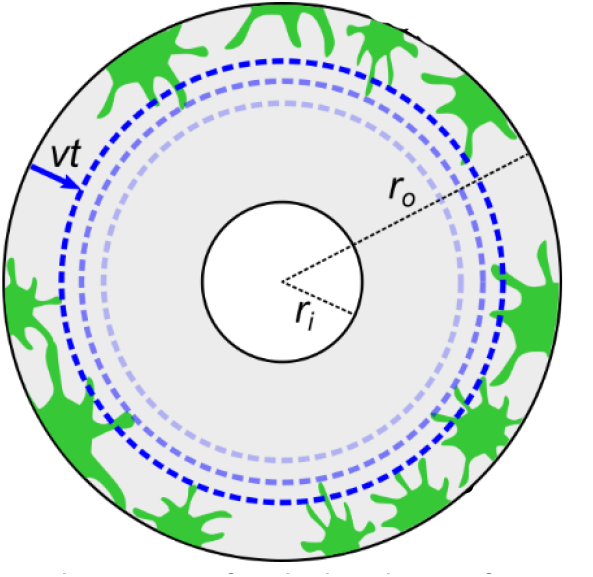

### Middle cerebral artery occlusion (MCAo)

To assess the functional contribution of microglial NKCC1 to ischemic brain injury, mice were subjected to a 45 min long MCAo (Middle Cerebral Artery occlusion) using a silicone-coated filament, as described earlier by Dénes et al. [85]. Surgery was performed under isoflurane (1,5% in a 30% O_2_ and 70% N_2_O gas mixture) anesthesia and core body temperature was tightly controlled (37 °C ± 0.5 °C) during the whole procedure using a homeothermic heating blanket. Laser Doppler flowmetry was used to validate the occlusion of the MCA. In brief, after a midline incision made on the ventral surface of the neck, the right common carotid artery (CCA) was isolated and a silicone-coated monofilament (210-230 μm tip diameter, Doccol, Sharon, US) was introduced to the left external carotid artery (ECA) and advanced along the internal carotid artery (ICA) to occlude the MCA. Animals were kept in a post-operative chamber at 26-28 °C until the functional assessment having free access to mashed food and water. Neurological outcome was assessed at 24h reperfusion using corner test and the 5-point Bederson’s sensory-motor deficit scoring system [86–88]. Briefly, the following scores were given: a 0, no motor deficit; 1, flexion of torso and contralateral forelimb when mouse was lifted by the tail; 2, circling to the contralateral side when mouse is held by the tail on a flat surface, but normal posture at rest; 3, leaning to the contralateral side at rest, 4, no spontaneous motor activity, 5, early death due to stroke. Functional outcome has also been assessed by a complex neurological scoring system to obtain a more comprehensive readout [89]. Results are expressed as composite neurological score. Composite scores range from 0 (healthy mice) to 56 (worst performance) by adding up scores from 13 categories as follows: hair (0–2), ears (0–2), eyes (0–4), posture (0–4), spontaneous activity (0–4), and epileptic behavior (0–12), and focal deficits: body symmetry (0–4), gait (0–4), climbing on a surface held at 45° (0–4), circling behavior (0–4), front limb symmetry (0–4), compulsory circling (0–4), and whisker response to a light touch (0–4).

Infarct volume and brain edema were calculated after 24h survival on cresyl violet stained coronal brain sections using ImageJ as described previously [90]. In brief, lesion volume was determined at eight neuro-anatomically defined coronal levels (between +2 mm rostral and -4 mm caudal to bregma) by integrating measured areas of damage and correcting for edema size. The pre-determined inclusion criteria for analysis were as follows; decline in Doppler signal of at least 70%, no cerebral hemorrhages and survival to 24 h. Cerebral hemorrhage was identified post-mortem by the presence of excessive bleeding on the external surface of the brain, typically close to the filament location.

### Immunohistochemistry

#### Perfusion, tissue processing and immunostaining for histology

Mice were anesthetized and transcardially perfused with 0.9% NaCl solution for 1 minute, followed by 4% PFA in 0.1 M phosphate buffer (PB) for 40 minutes, followed by 0.1 M PB for 10 minutes to wash the fixative out. Blocks containing the primary somatosensory cortex and dorsal hippocampi were dissected, and coronal sections were prepared on a vibratome (VT1200S, Leica, Germany) at 50 μm thickness for immunofluorescent histological and 100 μm thickness for the automated morphological analysis. Sections were washed in 0.1 M PB, incubated in 10%- and 30%-sucrose containing 0.1 M PB for 3 and 12 hours, respectively. Then the samples were placed into cryovials, snap frozen in liquid nitrogen, and stored at -80 °C for further use. For the free-floating immunohistochemical detection of NKCC1 each vial contained sections from NKCC1^fl/fl^ (WT), NKCC1^fl/fl ΔCx3CR1^ (KO) and NKCC1^-/-^ mice, which sections were marked by different cuts, enabling that all experimental parameters were completely identical for all samples. The sections were washed in TBS, blocked with a Mouse-on-Mouse blocker solution (MOM blocker, #BMK-2202, Vectorlabs) for 1 hour, washed in TBS 2x5 minutes, washed with MOM-diluent 2x5 minutes. The diluted anti-NKCC1 primary antibodies (NKCC1 Rb: 1:4000, #13884-1-AP, Proteintech, NKCC1 M: 1:2000, diluted in MOM-diluent; DSHB) were pre-incubated for 48 hours with brain slices from NKCC1^-/-^ mice, in order to remove the fraction of immunoglobulins that could potentially cause aspecific-binding. After discarding these slices, the guinea pig anti-Iba1 antibody (#234004, Synaptic Systems) was added to reach a 1:1000 final dilution. This antibody-mixture was applied on the samples for 48 hours at 4 °C during gentle shaking. After intensive washes in TBS, sections were incubated with a mixture of secondary antibodies (donkey-anti mouse Alexa 647, 1:1000, #715-605-150; donkey-anti rabbit Alexa 488, 1:1000, #711-546-152; donkey-anti guinea pig Alexa 594, 1:1000, #706-586-148; Jackson ImmunoResearch, diluted in TBS) for 24 hours. After subsequent washes in TBS and 0.1 M PB, DAPI staining was performed and the sections were mounted on glass slides, and covered with Diamond Antifade (#P36961, ThermoFisher) or Aquamount (#18606-5, Polysciences). Immunofluorescence was analyzed using a Nikon Eclipse Ti-E inverted microscope (Nikon Instruments Europe B.V., Amsterdam, The Netherlands), with a CFI Plan Apochromat VC 60X oil immersion objective (numerical aperture: 1.4) and an A1R laser confocal system. We used 405, 488, 561 and 647 nm lasers (CVI Melles Griot), and scanning was done in line serial mode. Single image planes were exported from the ND2 files, the outline of microglial cells was drawn using the Iba1-labeling, and the integrated density of the NKCC1 fluorescence signal was measured within these respective ROI-s. This section is related to Figure 2.

#### Perfusion, processing and immunostaining of ischaemic and LPS-injected tissues

In terminal ketamine-xylazine anesthesia (100 mg/kg-10 mg/kg) mice were transcardially perfused with 0.9% NaCl solution, followed by 4% phosphate buffered PFA. Brains of microglial NKCC1 KO mice subjected to 45 min MCAo were postfixed and cryoprotected overnight (in 4% phosphate buffered PFA-10% sucrose solution), then immersed into a cryoprotective solution (10% sucrose in PBS) at least 2h before 25 μm coronal sections were cut using a sledge microtome. Immunofluorescent staining was performed on free-floating coronal brain sections. Brain sections were blocked with 5% normal donkey serum followed by overnight incubation at 4 °C using the following mixture of primary antibodies: goat anti-IL-1β/ILF2 (1:250; #AF-401-NA, R&D Systems), rat anti-CD45 (1:250; #MCA1388, AbD Serotec), rabbit anti-P2Y12R (1:500; #55043AS, AnaSpec). After the incubation with the primaries, sections were washed several times in TBS and were incubated with mixture of corresponding secondary antibodies at room temperature for 2h. The following secondaries were used: donkey-anti goat CF568 (1:1000; #20106, Biotium), donkey-anti rabbit Alexa 647 (1:1000; #711-605-152, Jackson ImmunoResearch), donkey-anti rat Alexa 488 (1:1000; #712-546-153, Jackson ImmunoResearch). Slices were washed in TBS and were mounted to microscope slides using Fluoromount-G (#0100-01, SouthernBiotech). Representative images were captured with a 20X objective (Plan Apo VC, numerical aperture: 0,.75, FOV=645.12um) on a Nikon A1R confocal system. Quantitative analysis was performed on widefield images, captured with a 20X objective (Plan, numerical aperture: 0.4) on a Nikon Eclipse Ti-E inverted microscope (Nikon Instruments Europe B.V., Amsterdam, The Netherlands). IL-1α and IL-1β positive cells in the penumbral region, P2Y12R positive and CD45 positive cells in the whole cortex were counted on 3-3 serial coronal sections for a given brain area. This methodical description is related to Figure 4.

The following primaries and secondaries were used for GFAP and AQP4 immunolabeling: chicken anti-GFAP (1:1000, #173006, Synaptic Systems) and guinea pig anti-AQP4 (1:500, #429004, Synaptic Systems), donkey anti-chicken A594 (1:500, #703-586-155, Jackson ImmunoResearch) and donkey anti-guinea pig A647 (1:500, #706-606-148, Jackson ImmunoResearch). For imaging, fluorescent slide scanner (Panoramic MIDI 3D HISTECH) with 20X Plan-Apochromat objective was used. Raw integrated densities were automatically measured on all images in selected ROIs from the striatum, and then their per-animal average was calculated and used for statistical analysis. This methodical description is related to Supplementary Figure 2 and 5.

### Electrophysiology

#### Slice preparation

In all cases, mice (56-65 days) were decapitated under deep isoflurane anaesthesia. The brain was removed and placed into an ice-cold cutting solution, which had been bubbled with 95% O_2_–5% CO_2_ (carbogen gas) for at least 30 min before use. The cutting solution contained the following (in mM): 205 sucrose, 2.5 KCl, 26 NaHCO_3_, 0.5 CaCl_2_, 5 MgCl_2_, 1.25 NaH_2_PO_4_, 10 glucose, saturated with 95% O_2_– 5% CO_2_. Horizontal hippocampal slices of 250 µm thickness were cut using a Vibratome (Leica VT1000S). Slices were placed into an interface-type incubation chamber which contained standard ACSF at 35°C that gradually cooled down to room temperature. At the beginning of the incubation period, a solution of 25µm/ml Alexa 594 conjugated isolectin B4 (I21413, ThermoFisher) diluted in ACSF was pipetted on top of each slice (15 ml/slice). After this, slice preparations were incubated in darkness for 1 hour before measurement. All measurements took place within 4 hours after slicing. The ACSF solution contained the following (in mM): 126 NaCl, 2.5 KCl, 26 NaHCO_3_, 2 CaCl_2_, 2 MgCl_2_, 1.25 NaH_2_PO4, 10 glucose, saturated with 95% O_2_–5% CO_2_.

#### Recording conditions

After incubation for at least 1 hour, slices were transferred individually into a submerged-type recording chamber with a superfusion system allowing constantly bubbled (95% O_2_–5% CO_2_) ACSF to flow at a rate of 3-3.5 ml/min. The ACSF solution in all cases was adjusted to 300-305 mOsm to have a consistent normotonic medium during experiments, and in the hypotonic condition the same solution was diluted to have 50% the osmolarity of normotonic solution. Both normotonic and hypotonic ACSF solutions were constantly saturated with 95% O_2_–5% CO_2_ during measurements. All measurements were carried out at room temperature. A stock solution of 100 mg/ml gramicidin B (G5002, Sigma-Aldrich) diluted in DMSO was prepared daily, and further diluted to 100 µg/ml concentration in filtered, intracellular solution. Alexa 488 (100 µM) was also added to monitor membrane integrity during measurements. Before each recording, standard patch pipettes were fabricated from borosilicate glass and prefilled with gramicidin free intracellular solution, and then backfilled with the intracellular solution containing gramicidin and Alexa 488. The intracellular solution contained the following (in mM): 120 KCl, 1 CaCl_2_, 2 MgCl_2_, 10 HEPES, and 11 EGTA, pH: 7.3, 280-300 mOsm. Standard patch electrodes were used in all of the recordings, pipette resistances were 3-6 MΩ when filled with intracellular solution. Visualization of slices and selection of cells (guided by isolectin B signal) were carried out under an upright microscope (BX61WI; Olympus, Tokyo, Japan equipped with infrared-differential interference contrast optics and a UV lamp). Only cells below 15-20 µm measured from slice surface were targeted. During recordings, Alexa 488 signal was constantly monitored, to make sure that the membrane of cells did not suffer a rupture. Those cells who developed Alexa 488 signal in their cell body, were automatically discarded. All cells were recorded in voltage-clamp mode at -40 mV holding potential. Series resistance was constantly monitored and perforation was considered to be formed when values decreased to the 35-50 MΩ range (this regularly happened after 25-30 minutes after Gigaseal formation), and showed stability in series resistance and current response values between a 15% margin during the whole recording. Cells exceeding their initial value of series resistance and current responses measured at the beginning of both normotonic and hypotonic conditions beyond 15% in either directions during recordings were discarded. Input resistance of cells were 4.75 GΩ ± 1.56 SEM in WT and 3.94 GΩ ± 0.97 SEM in NKCC1 KO animals. There was no significant difference in input resistance values of measured cells between WT and NKCC1 KO animals (independent t-test, p=0.33). A pulse train ranging from -100 mV to 100 mV with 20 mV increments were recorded, each stimulus was 100 ms in length, and repeated 3 times. Gaps between pulses were 2000 ms in length. Pulse trains were recorded in the normotonic ACSF when stable perforated patch-clamp mode was established, and after 5 minutes of perfusion with hypotonic ACSF (50% dilution). Recordings were performed with a Multiclamp 700B amplifier (Molecular Devices). Data were digitized at 10 kHz with a DAQ board (National Instruments, USB-6353) and recorded with a custom software developed in C#.NET and VB.NET in the laboratory. Analysis was done using custom software developed in Delphi and Python environments.

#### Automated morphological analysis of microglial cells

100 µm thick mouse brain coronal sections were immunostained with guinea pig anti-Iba1 (1:500; #234004, Synaptic Systems), Alexa 647 donkey anti-guinea pig (1:500; #706-606-148, Jackson ImmunoResearch) antibodies, and DAPI. Imaging was carried out in 0.1M PB, using a Nikon Eclipse Ti-E inverted microscope (Nikon Instruments Europe B.V., Amsterdam, The Netherlands), with a CFI Plan Apochromat VC 60X water immersion objective (numerical aperture: 1.2) and an A1R laser confocal system. For 3-dimensional morphological analysis of microglial cells, the open-source MATLAB-based Microglia Morphology Quantification Tool was used (available at https://github.com/isdneuroimaging/mmqt). This method uses microglia and cell nuclei labeling to identify microglial cells. Briefly, 59 possible parameters describing microglial morphology are determined through the following automated steps: identification of microglia (nucleus, soma, branches) and background, creation of 3D skeletons, watershed segmentation and segregation of individual cells [47].

#### Quantification and Statistical analysis

All quantitative assessment was performed in a blinded manner. Based on the type and distribution of data populations (examined with Shapiro-Wilk normality tests) we applied appropriate statistical tests: In the case of two independent groups, Student’s t-test or Mann-Whitney U-test, for three or more independent groups one-way ANOVA followed by Tukey’s *post hoc* comparison or Kruskal-Wallis test with Dunn’s multiple comparison test was applied. Data were analyzed using the GraphPad Prism version 8.2 for Windows software (GraphPad Software, San Diego, California USA). In this study, data are expressed as mean±SEM, p<0.05 was considered statistically significant.

## Acknowledgments

This work was supported by „Momentum” research grant from the Hungarian Academy of Sciences (LP2016-4/2016 to A.D.) and ERC-CoG 724994 (to A.D.). Additionally, this work was funded by Hungarian National Scientific Research Fund (NKFIH-OTKA Grant No. K131844 to S.B.), the János Bolyai Research Scholarship of the Hungarian Academy of Sciences (to N.L. and C.C.), ÚNKP-20-3-II (to B.P.) and ÚNKP-20-5 (to C.C.) of the New National Excellence Program of the Ministry for Innovation and Technology, Hungary; and German Research Foundation (SPP 1665) and the Federal Ministry of Education and Research (NEURON ACRoBAT) to C.A.H. A.A. holds a Stipendium Hungaricum Scholarship from the Government of Hungary. We thank László Barna and the Nikon Imaging Center at the Institute of Experimental Medicine for kindly providing microscopy support, and thank Dóra Gali-Györkei for her excellent technical assistance.

## Author Contributions

Experimental design and overall concept, A.D. and Zs.K; Methodology, C.A.H.; N.L.; E.S.; P.B.; A.G.; Formal Analysis K.T.; N.L., B.P., S.B.; P.B.; D.K.; Investigation, K.T., N.L., B.P., R.F., E.S., A.A., C.C, Zs.K.; P.B.; Resources, A.D., S.B., C.A.H; Writing – Original Draft, K.T., Zs.K., A.D., Writing – Review & Editing K.T.; Zs.K., K.K., A.D. with all authors; Intellectual contribution, conceptual support K.K., C.A.H; Visualization K.T., E.S., C.C., B.P. D.K.; Supervision A.D. and Zs.K.; Project Administration A.D. and Zs.K.; Funding Acquisition, A.D.

