## Supplementary Figures for "NKCC1 modulates microglial phenotype, cerebral inflammatory responses and brain injury in a cell-autonomous manner"

### Supplemental Material

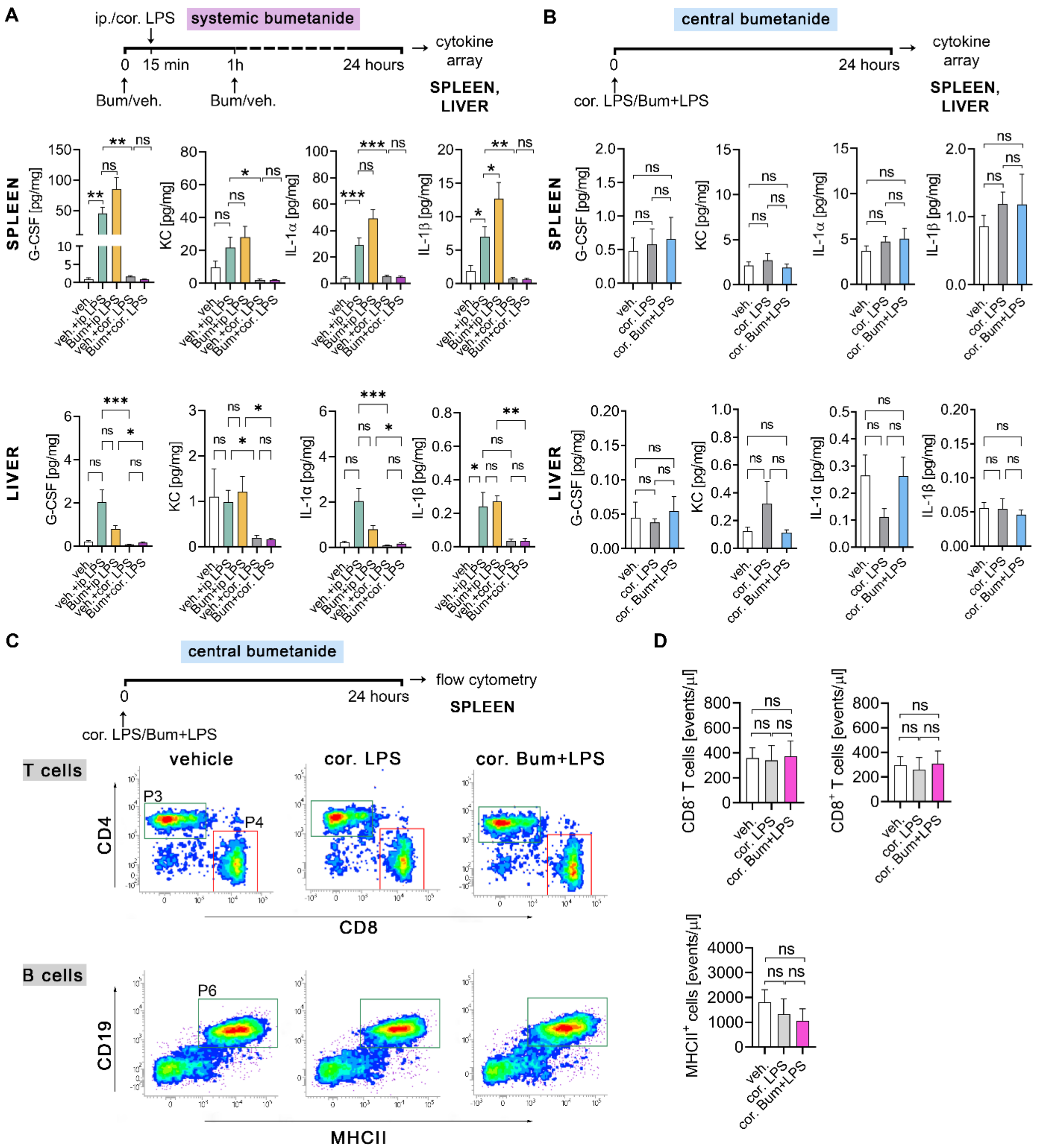


**Supplementary Figure 1. Effects of systemic vs. central blockade of NKCC1 on LPS-induced cytokine responses in the periphery. A**: Intraperitoneal bumetanide (Bum) injections further enhance the ip. LPS-induced GCSF, IL-1α and IL-1β production, while intracortical LPS alone or with Bum has no significant effect on cytokine levels in the spleen and liver. **B:** Intracortical (cor.) LPS injection does not induce cytokine production and Bum has no effect on baseline cytokine levels in the spleen and liver. **C-D:** Flow cytometric dot plots show that cortical administration of Bum does not alter the numbers of $CD4^+$ (P3 gate) and $CD8^+$ (P4 gate) T cells and $CD19^+$ $MHCII^+$ (P6 gate) B cells in the spleen. All data are expressed as mean±SEM. **A-B**: One-way ANOVA followed by Tukey's multiple comparison test; *p<0.05; **p<0.01; ***p<0.001; **A:** N=6/group; **B:** N=9/group. **D:** One-way ANOVA followed by Tukey's multiple comparison test; N=4/group Abbreviations: veh.: vehicle; ip: intraperitoneal; cor.: cortical; Bum: bumetanide; ns: not significant.

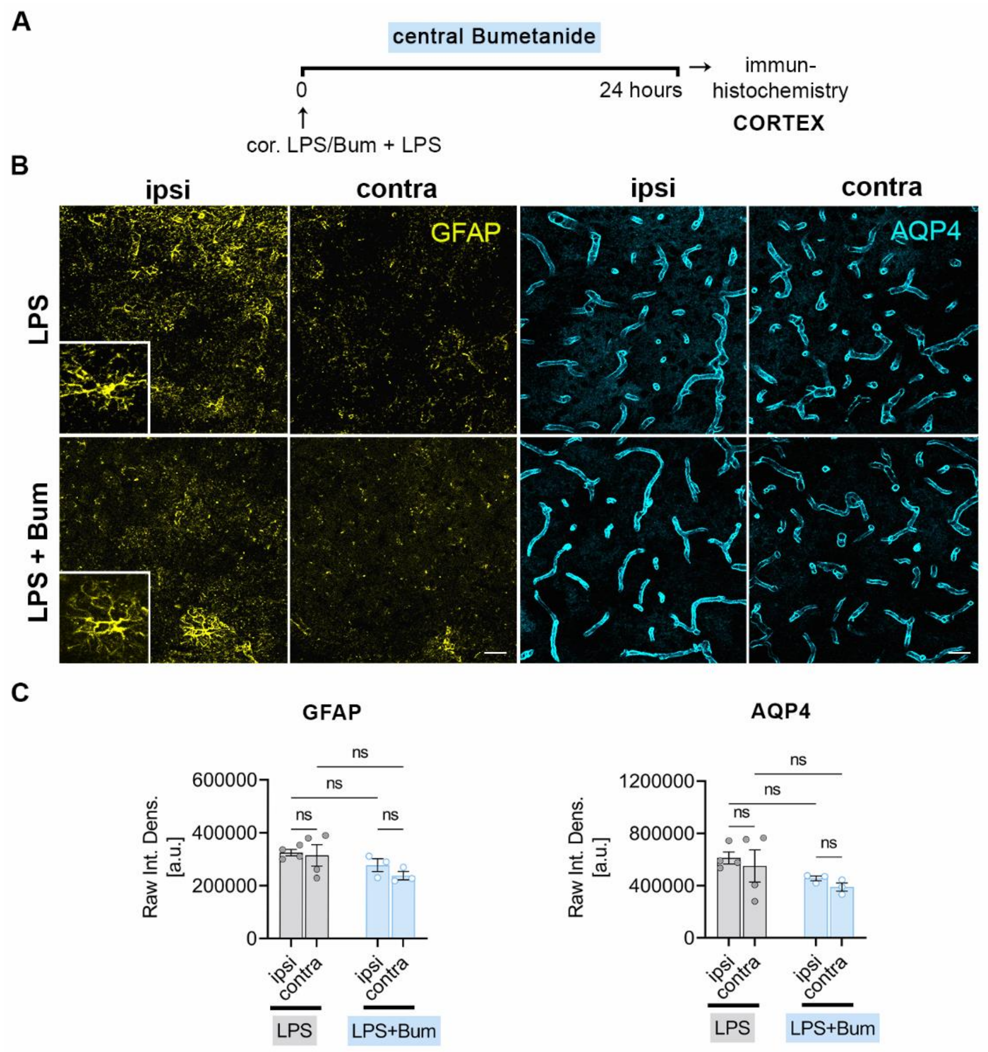


**Supplementary Figure 2. Intracortical blockade of NKCC1 does not alter astroglial GFAP or AQP4 levels. A-B:** CLSM images show immunolabeling for GFAP (yellow) and AQP4 (cyan) in NKCC1$^{fl/fl}$ animals 24 hours after cortical injection of LPS or LPS+Bum and in the corresponding contralateral areas. **C:** Raw integrated densities were automatically measured on all images in randomly selected ROIs from the injected ipsilateral cortical and contralateral regions prior to statistical analysis. No statistically significant difference in GFAP and AQP4 expression levels is seen in parenchymal astrocytes or perivascular astrocyte endfeet. B: Scale: 25 µm **C:** One-way ANOVA followed by Holm-Sidak's multiple comparisons test; N=4 mice/group; Abbreviations: ns: not significant, Bum: bumetanide

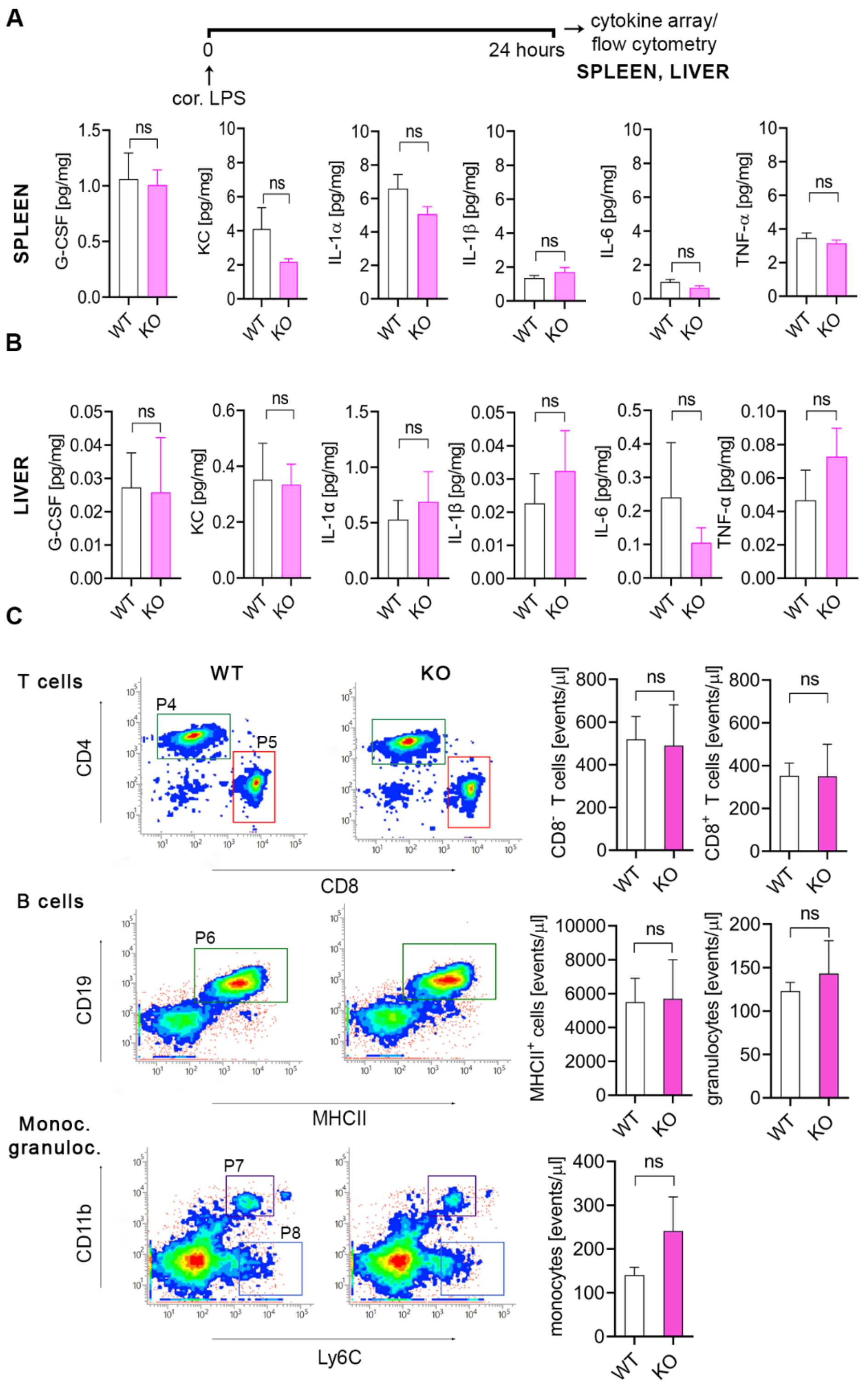


**Supplementary Figure 3. Microglial NKCC1 deficiency does not alter cytokine levels and main leukocyte populations in the spleen after intracortical LPS injection. A-B:** Cytokine levels in the spleen and liver do not differ between WT and NKCC1 KO mice after intracortical LPS administration. **C:** Numbers of $CD4^+$ (P4 gate), $CD8^+$ (P5 gate) T cells, and $CD19^+$ $MHCII^+$ B cells (P6 gate) are not altered in the spleen of NKCC1 KO mice compared to WT. Microglial NKCC1 deficiency does not affect the proportion of monocytes (P8 gate) or granulocytes (P7 gate) compared to WT. **A-B:** Mann-Whitney test, *: *p* <0.05; N (WT)=7, N (KO) =6; **C:** Unpaired t-test; N (WT)=4, N (KO)=4; n. s.: not significant.

A

B

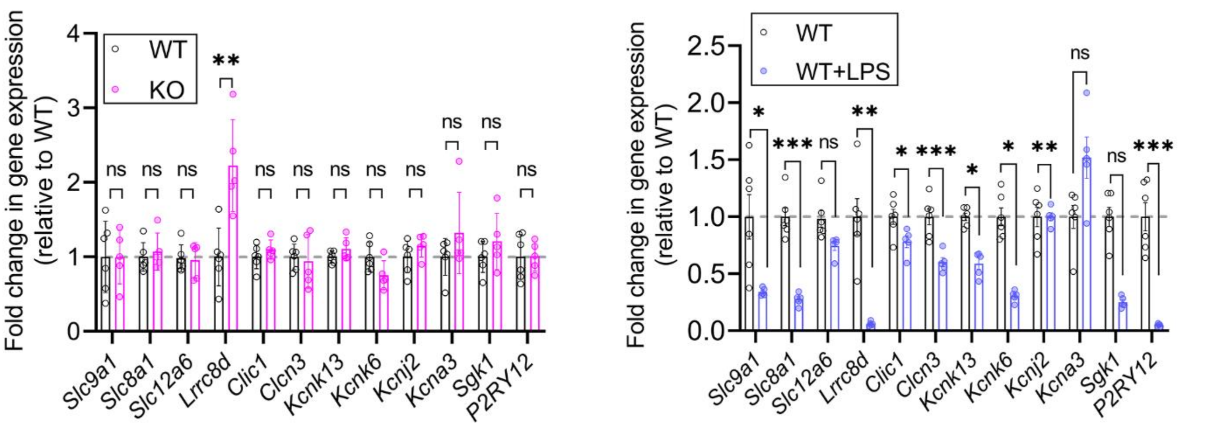


C

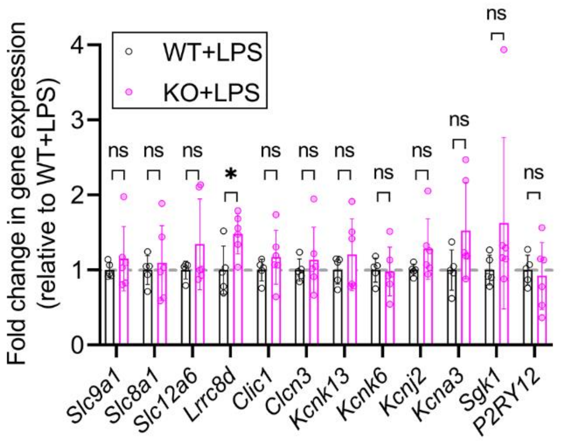


D

| Gene symbol | Description | Reference(s) |
|---|---|---|
| ***Slc9a1*** | NH1E, Na$^+$/H$^+$ exchanger | (39, 56) |
| ***Slc8a1*** | NCX1, Na$^+$/Ca$^{2+}$ exchanger | (39, 56) |
| ***Slc12a6*** | KCC3, chloride transmembrane transport | (2, 39) |
| ***Lrrc8d*** | VRAC subunit D, volume-sensitive anion channel activity | (39, 56) |
| ***Clic1*** | chloride intracellular channel 1, plasma membrane chloride conductance | (39, 56) |
| ***Clcn3*** | voltage-gated chloride channel activity | (39) |
| ***Kcnk13*** | THIK-1, two pore domain potassium channel | (34, 57) |
| ***Kcnk6*** | TWIK-2, two pore domain potassium channel | (34) |
| ***Kcnj2*** | Kir.2.1, inwardly-rectifying potassium channel activity | (35, 56) |
| ***Kcna3*** | Kv1.3, voltage-gated potassium channel activity | (35, 56) |
| ***Sgk1*** | serine-threonine kinase, cell volume regulation | (39, 48) |
| ***P2RY12*** | G-protein coupled purinergic receptor | (34, 57) |

**Supplementary Figure 4. Changes in mRNA levels of microglial ion channels, transporters, and exchangers in the absence of microglial NKCC1 and after LPS treatment. A:** The expression of most genes that contribute to ion regulation, membrane potential and cell volume regulation (anion channels (CLIC1); K$^+$ channels (Kv1.3; Kir2.1; THIK-1; TWIK-2); ion exchangers (NH1E Na$^+$/H$^+$ exchanger; NCX1 Na$^+$/Ca$^{++}$ exchanger; CLCN3 H$^+$/Cl$^-$ exchanger), transporters (KCC3 K$^+$/Cl$^-$ transporter) are not altered in NKCC1 KO microglia. However, Lrrc8d mRNA levels show a two-fold increase in NKCC1 KO microglia cells. **B:** *Slc9a1*, *Slc8a1*, *Lrrc8d*, *Clic1*, *Clcn3*, *Kcnk13*, *Kcnk6*, *Kcnj2*, *Sgk1* gene show decreased expression level in microglial cells 24 hours after intracisternal LPS injection. **D:** Summary table of investigated genes. **A-C:** Unpaired t-test, **A:** N (WT)=6, N (KO)=5; **: $p$<0.01, **B:** N (WT)=6, N (WT+LPS)=5, *: $p$<0.05, **: $p$<0.01, *** $p$<0.001, **C:** N (WT+LPS)=5, N (KO+LPS)=6, *: $p$<0.05; ns: not significant

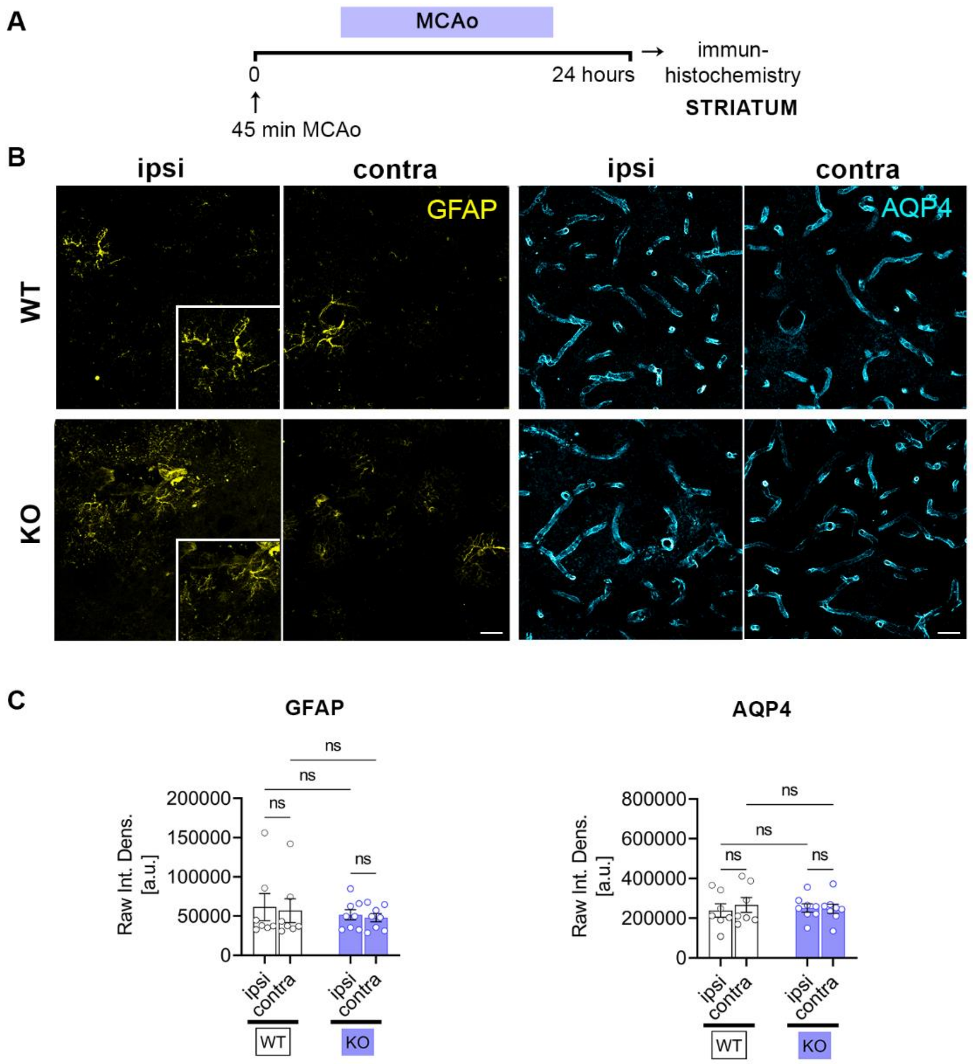


**Supplementary Figure 5. Deletion of microglial NKCC1 does not alter astroglial GFAP and AQP4 levels. A-B:** CLSM images show immunolabeling for GFAP (yellow) and AQP4 (cyan) in microglial NKCC1 KO animals 24 hours after MCAo. **C:** Raw integrated densities were automatically measured on all images in selected ROIs from the striatum, then, their per-animal average was calculated and used for statistical analysis. Data show no statistically significant differences in GFAP and AQP4 expression levels. **C:** One-way ANOVA followed by Holm-Sidak's multiple comparisons tests; N (WT)=7, N (KO)=8 mice; Abbreviations: ns: not significant

**Supplementary Video 1.** *In vivo* 2P time-lapse imaging of Cx3CR1$^{+/GFP}$ mice shows microglial responses to focal lesion-induced injury under control conditions (left) and after cisterna magna Bumetanide injection (right). Lesion-induced microglial process recruitment was determined by model fitting using image data from the circular region marked on the video. The outer perimeter of this region corresponds to the lesion site.
